# Proteomic network analysis of bronchoalveolar lavage fluid in ex-smokers to discover implicated protein targets and novel drug treatments for chronic obstructive pulmonary disease

**DOI:** 10.1101/2022.02.14.480388

**Authors:** Manoj J. Mammen, Chengjian Tu, Matthew C. Morris, Spencer Richman, William Mangione, Zackary Falls, Jun Qu, Gordon Broderick, Sanjay Sethi, Ram Samudrala

## Abstract

**Rationale:** Bronchoalveolar lavage of the epithelial lining fluid can sample the profound changes in the airway lumen milieu prevalent in Chronic Obstructive Pulmonary Disease (COPD). Characterizing the proteins in bronchoalveolar lavage fluid in COPD with advanced proteomic methods will identify disease-related changes, provide insight into pathogenetic mechanisms and potential therapeutics that will aid in the discovery of more effective therapeutics for COPD.

**Objectives:** We compared epithelial lining fluid proteome of ex-smokers with moderate COPD who are not in exacerbation status COPD, to non-smoking healthy control subjects using advanced proteomics methods and applied proteome-scale translational bioinformatics approaches to identify potential therapeutic protein targets and drugs that modulate these proteins towards the treatment of COPD.

**Methods:** Proteomic profiles of bronchalveolar lavage fluid were obtained from 1) never-smoker control subjects with normal lung function (n=10) or 2) individuals with stable moderate (GOLD stage 2, FEV1 50% – 80% predicted) COPD who were ex-smokers for at least one year (n=10). NIH’s Database for Annotation, Visualization and Integrated Discovery (DAVID) and Ingenuity’s Ingenuity Pathway Analysis (IPA) were the two bioinformatics tools employed for network analysis on the differentially expressed proteins to identify potential crucial hub proteins. The drug-proteome interaction signature comparison and ranking approach implemented in the Computational Analysis of Novel Drug Opportunities (CANDO) platform for multiscale therapeutic discovery was utilized to identify potential repurposable drugs for the treatment of COPD based on the BALF proteome. Subsequently, a literature-based knowledge graph was utilized to rank combinations of drugs that would most likely ameloriate inflammatory processes by inhibition or activation of their functions.

**Results:** Proteomic network analysis demonstrated that 233 of the >1800 proteins identified in the BALF were differentially expressed in COPD versus control, including proteins associated with inflammation, structural elements, and energy metabolism. Functional annotation of the differentially expressed proteins by their implicated biological processes, cellular localization, and transcription factor interactions was accomplished via DAVID. Canonical pathways containing the differential expressed proteins were detailed via the Ingenuity Pathway Analysis application. Topological network analysis demonstrated that four proteins act as central node proteins in the inflammatory pathways in COPD. The CANDO multiscale drug discovery platform was used to analyze the behavioral similarity between the interaction signatures of all FDA-approved drugs and the identified BALF proteins. The drugs with the signatures most similar interaction signatures to approved COPD drugs were extracted with the CANDO platform. The analysis revealed 189 drugs that putatively target the proteins implicated in COPD. The putative COPD drugs that were identified using CANDO were subsequently analyzed using a knowledge based technique to identify an optimal two drug combination that had the most appropriate effect on the central node proteins.

**Conclusion:** Analysis of the BALF proteome revealed novel differentially expressed proteins in the epithelial lining fluid that elucidate COPD pathogenesis. Network analyses identified critical targets that have critical roles in modulating COPD pathogenesis, for which we identified several drugs that could be repurposed to treat COPD using a multiscale shotgun drug discovery approach.

## Introduction

Chronic obstructive pulmonary disease (COPD) is a leading cause of mortality and morbidity in the US.^1-5^ Additionally, COPD results in millions of hospitalizations in the developing world.^1,6-11^ The prevalence of cigarette smoking continues to rise in most developing countries around the world.^12-14^ However, only 25-50% of tobacco smokers develop COPD, suggesting only a subset develops an exaggerated inflammatory process that leads to lung destruction.^12,13,15^ Bronchoalveolar lavage fluid (BALF) and bronchial samples from ex-smokers reveal active inflammation long after smoking cessation.^16^

Although structural changes in the airways, parenchyma, and pulmonary vessels are typical in patients with COPD, the lower airways and the alveoli are the initial sites of the inflammatory process.^17,18^ The inflammatory process initiated by smoking persists after cessation and is likely exaggerated by autoimmunity and infection.^19,20^ Accurate and precise measurement of the molecular mediators in the airways should aid in rigorous analysis of their role in disease.

There has been a keen interest in understanding the genetic determinants of COPD, as the interaction between genes and environment leads to protein expression, ultimately resulting in either healthy or disease states. However, genomic data alone does not predict protein abundance or activity; proteins are the ultimate participants in integrated biological processes that lead to healthy physiological function or pathology. Proteome-based analysis of bronchoalveolar lavage fluid (BALF) in COPD can identify tissue-specific markers of inflammation that can lead to understanding the mechanisms of COPD progression.

We sought to determine an unbiased proteome-based analysis of BALF in COPD under stable conditions (not in exacerbation status) to identify a broad series of molecules involved in COPD pathogenesis. A label-free proteomics mass spectroscopy method was utilized. The differentially expressed proteins were analyzed using multiple bioinformatics tools to critical pathways that were altered in these ex-smoker patients with COPD compared to healthy, never smoker controls, proteins implicated in COPD etiology, and to identify putative drug candidates that can be repurposed to treat COPD.

The raw proteomic data used in this manuscript was initially detailed in a previously published methodology manuscript using strict criteria (2 peptide identification criteria for a protein, ≥1.5 fold change, and p-value<0.05) to identify 423 individual proteins with 76 proteins expressed differently between COPD and controls.^21^ In this analysis, we adopted a pragmatic approach to the same raw proteomic data (1 peptide identification criterion, ≥1.5 fold change, and p-value<0.05) that identified 1831 individual proteins and 233 differentially expressed proteins between the two groups. The latter, more practical, approach provides important additional information for biomarker and therapeutic target discovery that may be utilized in future research to discover useful interventions.

## Methods

We analyzed the protein quantifications derived from the BALF of subjects with COPD and healthy ex-smoker control subjects via liquid chromatography and mass spectroscopy. We then used pathway analysis tools to identify relevant cellular pathways associated with differentially expressed proteins quantified from the BALF analysis. We subsequently employed the Computational Analysis of Novel Drug Opportunities (CANDO) platform to identify FDA approved drugs that could be repurposed to COPD, based on their putative interaction with the differentially expressed proteins. Using topological network analysis, we identified putative hub proteins that modulate the cellular pathways associated with COPD. Using the medical literature to predict the repurposed drugs effects on the most important hub protein, we created a refined list of drugs predicted to modulate the cellular pathway in order to impede COPD pathogenesis. to generate proteomic interaction signatures for the compounds

### Recruitment of subjects

BALF was obtained in a NHLBI funded study of innate lung defense in COPD.^22^ All procedures received approval from the Institutional Review Board (IRB), Veterans Affairs Western New York Healthcare System (WNY-VA), and strictly adhered to institutional guidelines.

### Ethics statement

This study is a sub-study of a larger group of patients with COPD and healthy controls to understand biological determinants of exacerbation frequency and was approved by the Institutional Review Boards of the Veterans Affairs Western New York Healthcare System and University at Buffalo. The participants gave written consent to the study via an IRB-approved consent form.

### Inclusion/exclusion criteria

The inclusion criteria and procedures for this study have been described previously and are provided in the supplementary material. ^22^ After informed consent, 116 volunteers were divided into three groups: 1) healthy nonsmokers, 2) ex-smokers with COPD, and 3) active smokers with COPD and underwent bronchoscopy and bronchoalveolar lavage. The methodology for bronchoscopy, lavage, and sample processing is included in the supplementary material.

For this study, we selected BALF obtained from ten ex-smokers with moderate COPD and ten healthy non-smoking controls for proteomic analysis, respectively. To minimize variability due to effects of acute smoking and disease severity, we confined this analysis to ex-smokers and moderate stage 2 disease per the Global Obstructive Lung Disease (GOLD)^23^ criteria of the forced expiratory volume in 1 second (FEV1) 50-80% predicted. All ex-smokers had ceased smoking for at least one year.

### Bronchoscopy and BALF sample preparation

The research bronchoscopy and BALF sample preparation were performed as described previously.^24^

### Protein identification/quantification

To investigate the soluble molecules in the epithelial lining fluid that may participate in COPD pathogenesis, unbiased proteomic analysis of BALF commenced without protein depletion or fractionation. Details of the methodology have been published^25^ and are also provided in the supplementary material.

### Long gradient nano-RPLC/mass spectrometry

Complete separation of the complex peptide mixture utilized a nano-LC/nanospray setup;^26^ the ion-current long gradient approach with mass spectrometry and subsequent protein identification was performed as described in Tu, et al. ^25-27^ All proteins identified with one or more peptide hits, fold change of ≥1.5, and p-value <0.05 are included as part of the differentially expressed BALF proteome.

### Bioinformatics analyses

#### Manually curated pathway analysis

Gene ontology, transcription factors, and expression locations were determined by uploading the protein expression dataset onto a web-based tool, the NIH’s Database for Annotation, Visualization and Integrated Discovery (DAVID) v6.7 (http://david.abcc.ncifcrf.gov/). ^28,29^

Biological networks were generated with Ingenuity Pathway Analysis (IPA, Ingenuity Systems), a web-based relational database and network generator. Proteins overrepresented in the uploaded datasets in biological networks, canonical pathways, and biological processes were identified.

#### Literature informed protein-protein and protein-drug interaction network

In addition to annotating differentially expressed proteins with the manually curated pathways cataloged in IPA, a network of protein-protein interactions was created using known regulatory relationships extracted from published scientific literature using the MedScan text-mining engine^30^ as well as protein-drug interactions cataloged in the Reaxsys medicinal chemistry database (Elsevier, Amsterdam). These are embedded in the broader Elsevier Knowledge Graph database ^31^ and were accessed via the Pathway Studio interface (Elsevier, Amsterdam).^32^

#### Shotgun multiscale drug discovery platform

We used the Computational Analysis of Novel Drug Opportunities (CANDO) platform^33-40^ to predict drugs that can be repurposed for the treatment of stable COPD. In CANDO, a compound/drug is considered to be potentially repurposable for an indication when it is found to have similar binding interactions with a specific proteome or library of proteins as a drug with known approval for the indication of interest.

In this study, we calculated the interaction scores between 2,450 United States Federal Drug Administration (FDA) approved drugs from the CANDO version 2.3 compound library and a curated human library of 8,385 proteins, including 5,316 solved X-ray crystallography structures and 3,069 computed protein structures modeled by I-TASSER^41,42^. The interaction scores were calculated using the bioanalytic docking (BANDOCK) protocol in the CANDO which utilizes predicted binding site information and chemical similarity to determine an interaction score that is a surrogate for the likelihood of interaction between a compound and protein.^33^ Binding sites were predicted for all human proteins using COACH ^43^, which uses the consensus of three complementary methods utilizing structure and sequence information to find similarity to solved structures in the Protein Data Bank (PDB).^44,45^ For each binding site predicted by COACH, a confidence score (PScore) and an associated co-crystallized ligand are output. The ligand is then compared to the query compound/drug using chemical fingerprinting methods, which enumerates the presence or absence of molecular substructures on the compound/drug. The Sorensen-Dice coefficient^46^ between the protein-ligand and compound/drug fingerprints (CScore) is also computed. The BANDOCK interaction score outputted for each compound-protein pair is the product of the Pscore and the Cscore.

For this analysis, we focused on the differentially expressed proteins in the BALF proteome (as described), and drugs used to treat COPD (“MESH:D029424”) (Table S1). We selected proteins in the CANDO human protein library that were also represented in the differentially expressed BALF proteome. We then used the CANDO platform to predict the top drug candidates that could be repurposed to treat COPD based on compound-proteome interaction signature similiarity to drugs currently approved/used to treat stable COPD. The protocol iterates through 34 known drugs used to treat stable COPD, counting the number of times drugs not associated with COPD show up in the top30 most similar compounds to the known treatments, then outputs the consensus predictions ranked by the number of times each compound appeared across all top30 lists. The similarity between a given drug and all other drugs in the library is determined by comparing their proteomic interaction signatures using the cosine similarity metric, where compounds with greater similarity scores rank stronger than those with low similarity. Thereby drugs that were most similar (in terms of interaction signatures) to multiple drugs used to treat COPD will be ranked highest.

#### Network topological analysis

Although not a complete descriptor, the topological location, and aspects of the connectivity linking a node to a broader biological network can inform the node’s function in mediating network behavior. Among the measures of a node’s importance or centrality, *betweenness* centrality has been used to describe how a node might serve as an important mediator of information flow in a regulatory network. In this work, *C*_*b*(*n*)_ for each node *n* of a network was calculated using the Brandes algorithm.^47^ The betweenness centrality of a node *n* reflects the amount of control that this node exerts over the interaction between communities of neighboring nodes in the network^48^ and can be computed as follows:

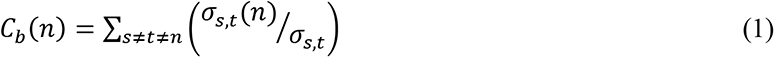

Where *s* and *t* are the source and target nodes in the network different from *n*, *σ*_*s,t*_ denotes the number of shortest paths from all *s* to all *t*, and *σ*_*s,t*_ is the number of shortest paths from s to *t* that must pass through node *n*. Here, unweighted betweenness centralities were calculated for each node in the literature-informed protein-protein network. The betweenness centrality scores for all nodes were expressed as fractions of the maximum betweenness centrality present in the network. All calculations were conducted in R version 4.0.2.^49^

#### Literature based drug enrichment analysis

Using putative drugs ranked by CANDO and further analyzed via the Elsevier Knowledge Graph,^31^ a drug enrichment analysis was performed to predict which drugs can most closely mimic an idealized intervention against the hub proteins identified in the network topological analysis. Drugs are represented as vectors with a length equal to the empirically derived number protein entities in the network model. Each index value is listed as 0 if there is no interaction between the drug and the corresponding model entity, a 1 if the drug promotes that entity, or a –1 if the drug inhibits that entity. Next, the cosine similarity, Sc, between each drug vector and the idealized intervention vector is calculated.^50^ Cosine similarity is calculated as:

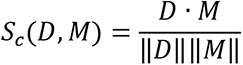

Where *D* is the drug vector and *M* is the idealized intervention. Higher Sc indicates a closer match between the drug vector and the idealized vector. A Sc of 1 means the two vectors are identical, and -1 indicates that the two are exactly opposed. For multidrug combinations, the net-effect of the individual drug vectors is calculated as:

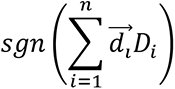

Where *n* is the total number of drugs in the combination, *Di* is the vector corresponding to the *i*th drug, and *sgn* is the sign function. The cosine similarity of the net-effect vector and idealized vector is then calculated.

The statistical significance of these enrichment scores is determined empirically from an estimated null distribution of cosine similarities. This null distribution uses a set of model-relevant background drugs for which each interacts with at least one entity in the network. All CANDO drugs of interest were included in the background. Empirical p-values are estimated as

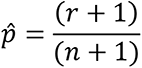

Where *r* is the number of null Sc values greater than the observed Sc and *n* is the total number of null Sc values.

### Statistical analysis

Statistical analysis was performed with SPSS/19. Demographic values were depicted as mean ± SEM.

## Results

### Study population characteristics

Characteristics for subjects included in the BALF study are shown in ***Table 1***, with the only significant differences between the two groups in tobacco smoke exposure and lung function.

**Table 1:** Clinical parameters of never-smoking healthy subjects and ex-smokers with stable COPD in BALF study FEV1: forced expiratory volume in 1 second. FVC: forced vital capacity. Years quit: Years subjects quit tobacco smoking. Pack-Years: The average number of packs of cigarettes smoked per week multiplied by the years the subject smoked cigarettes.

|  | Control subjects<br>n=10 | COPD subjects<br>(GOLD stage 2)<br>n=10 | P-value |
| --- | --- | --- | --- |
| **Age (years)** | 63.4 ± 11.7 | 67.8 ± 8.5 | 0.15 |
| **Sex** |  |  | 0.31 |
| Male | 6 | 7 |  |
| Female | 4 | 3 |  |
| **Race** |  |  | 0.083 |
| Caucasian | 8 | 10 |  |
| African-American | 2 | 0 |  |
| **BMI (kg/m<sup>2</sup>)** | 28.5±4.2 | 32±9.7 | 0.32 |
| **Years patient quit smoking** | NA | 12.9 ± 4.4 |  |
| **Tobacco smoking, Pack years** | NA | 56.6 ± 17.2 | <0.001 |
| **FEV<sub>1</sub> (% predicted)** | 96.3 ± 14.8 | 65.9 ± 8.1 | <0.001 |
| **FVC (% predicted)** | 95.6 ± 13.4 | 87.6± 13.1 | 0.19 |
| **FEV<sub>1</sub>/FVC** | 77.6± 3.8 | 57.8 ± 8.6 | <0.001 |

### BALF proteome characteristics

A total of 1831 unique proteins were identified in the BALF proteome. A total of 233 proteins (>1.5-fold absolute change, p-value <0.05) had a significant differential expression in BALF samples from patients with COPD versus healthy ex-smokers, 138 proteins were decreased in COPD while 95 proteins were increased (***Table S2 and Table S3*).**

### Manually curated pathway analysis

#### Functional annotation of differential expressed proteins and transcription factor interactions

The 233 differentially quantified proteins were characterized by their biological processes, transcription factor interactions, and cellular localization by employing NIH’s DAVID.^28,29^ The proteins involved in several biological processes implicated in COPD pathogenesis (total number of proteins, number upregulated, number downregulated) such as proteolysis^51^ (20,4,16), extracellular matrix^52^ (13,6,7), cell adhesion^53^ (11,2,9), cytoskeleton^54^ (32,14,18), defense response^55^ (16, 7,9), cell migration^56^ (12,4,8), and oxidation-reduction^21^ (11,2,9) were altered in COPD. As expected with examining the lung lining fluid, the largest single group of differentially expressed proteins was associated with the extracellular space (49, 30, 19).

Transcription factors (**Table S4**) associated with the differentially expressed proteins (total number of proteins associated with the transcription factor) included serum response factor-SRF (148), transcription factor 8-AREB6 (166), signal transducer and activator of transcription factor 1-STAT1 (69), zinc finger protein-GFI1 (97), signal transducer and activator of transcription factor 3-STAT3 (101), nuclear factor kappa-light-chain-enhancer of activated B cells-NF-**κ**B (79), CCAAT/enhancer-binding protein **β**-CEBPB(109), paired box gene 2-PAX2(113), and activating transcription factor 2-CREBP1(95).

#### Bioinformatic pathway analysis of BALF proteomic data

The protein expression datasets were imported into IPA (Ingenuity Systems) and projected onto the relevant biological pathways; processes linked to the differentially expressed proteins were analyzed with IPA’s manually curated knowledge database. Of the 233 differentially expressed proteins, 217 matched to the IPA curated database and were analyzed. Sixteen pathways were noted to have several proteins associated with the differentially expressed BALF dataset (**Table S5**), including proteins implicated in cellular movement, cellular death and survival, cell morphology, immune cell trafficking, and cell cycle. Appendix **Figures S1 to S4** depict IPA networks of selected pathways with the highest number of differentially expressed proteins.

### Computational drug prediction

130 out of 233 BALF differentially expressed proteins were identified in the CANDO human protein library. This subset of proteins within the CANDO platform was used to predict 189 putative drug candidates that have the most similar protein interaction signatures to the set of known drugs used to treat COPD ( **Figure 1** and **Table S6**). Many of the drugs were corticosteroids; however other putative drugs included tezacaftor^57^, a recently developed drug to potentiate sodium channel activity in the treatment of cystic fibrosis; two additional drugs predicted to treat COPD, gemfibrozil^58^, and pioglitazone^59^, are drugs currently used to treat hyperlipidemia and diabetes, respectively.

**Figure 1.**
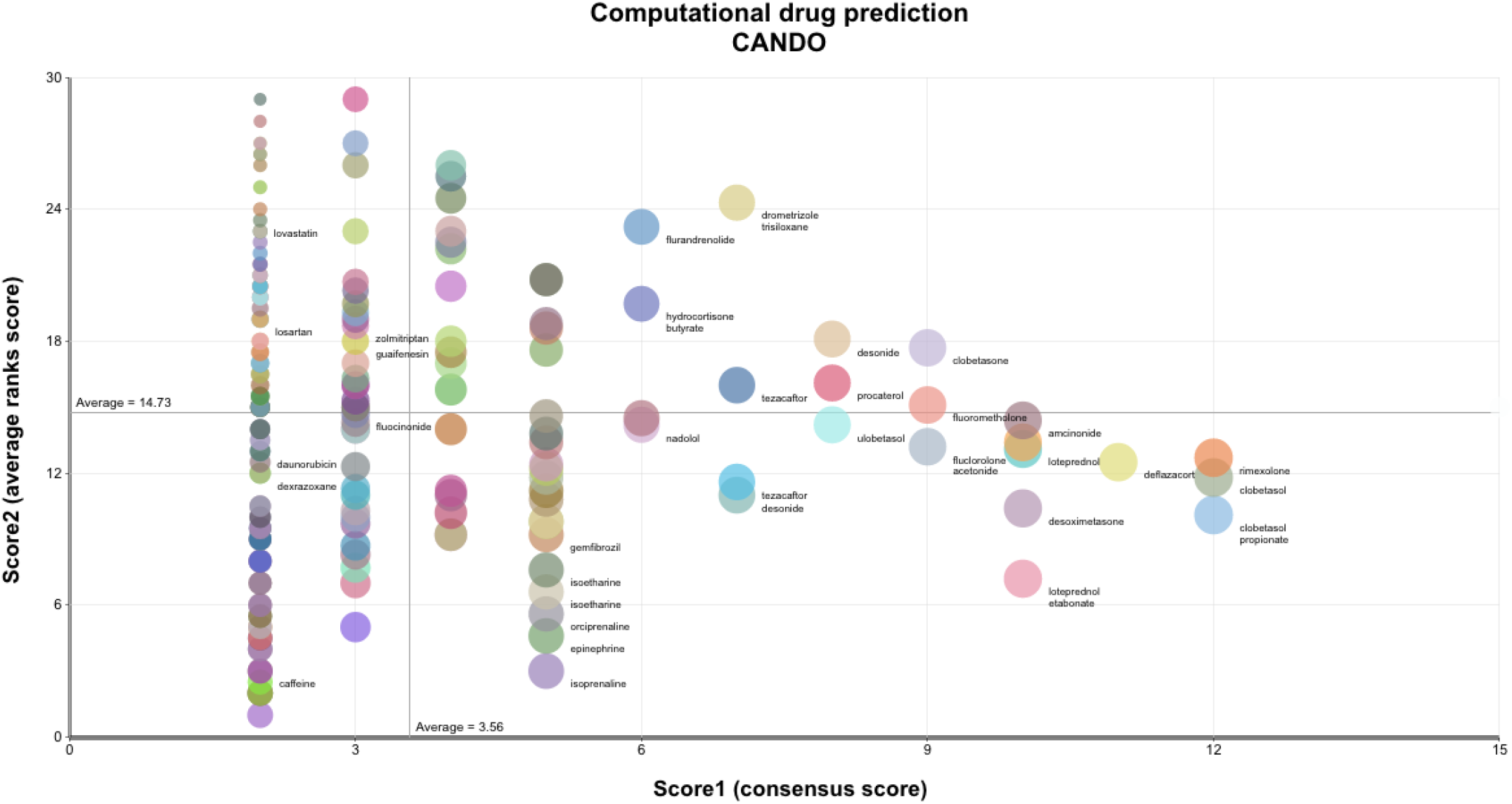
Putative drug candidates for treating COPD generated using the CANDO platform. A subset of 130 proteins from the CANDO human protein library were identified from 233 differentially expressed proteins in the BALF. These 130 proteins were utilized to generate BALF-specific interaction signatures for 2,450 FDA-approved drugs via our in-house docking protocol BANDOCK (see methods). These drug-proteome interaction signatures were compared to those of 34 known drugs used to treat COPD to predict 189 most similar putative drug candidates. The 189 drugs are represented by colored circles, with the diameter of the circles decreasing with descending overall rank. Drug name labels are depicted for a selection of the 189 drugs shown by the colored circles. The horizontal axis plots the consensus score count or the number of times the particular drug is listed within the top30 most similar drugs to those known to treat COPD based on interaction signature similarity. The vertical axis plots the average of the cumulative ranks of the consensus scores for the putative drug. The overall rank of a putative drug is determined by initially sorting the drug by the consensus score, as noted above, and then additional sorting by the average rank. Many of the drug candidates were corticosteroids not used to treat COPD; however other putative drugs included tezacaftor, a drug to potentiate sodium channel activity in the treatment of cystic fibrosis; two additional drugs predicted to treat COPD, gemfibrozil, and pioglitazone, are drugs currently used to treat hyperlipidemia and diabetes, respectively. This analysis indicates that the CANDO platform applied to the BALF proteome is able to generate putative drug candidates for COPD treatment. BANDOCK= bioanalytical docking CANDO=computational analysis of novel drug opportunities COPD=chronic obstructive pulmonary disease.

### Candidate key mediators of COPD pathology based on literature derived drug enrichment

#### Literature informed protein-protein and protein-drug interaction network

A total of 233 proteins were identified as differentially expressed between COPD patients and healthy controls by mass spectrometry. Of these, 214 were represented in the Elsevier Knowledge Graph^31^, with the remainder comprising specific immunoglobulin chain proteins. A query of the Knowledge Graph for documented regulatory interactions between these protein entities yielded 206 regulatory edges supported by 807 references (with a median of 1 reference per edge). 112 of the 214 identified proteins could not be connected to the broader network circuit by a documented interaction. The protein entities in this network were then assessed in terms of their importance as mediators of signal transfer based on their betweenness centrality. (**Figure 2**).

**Figure 2.**
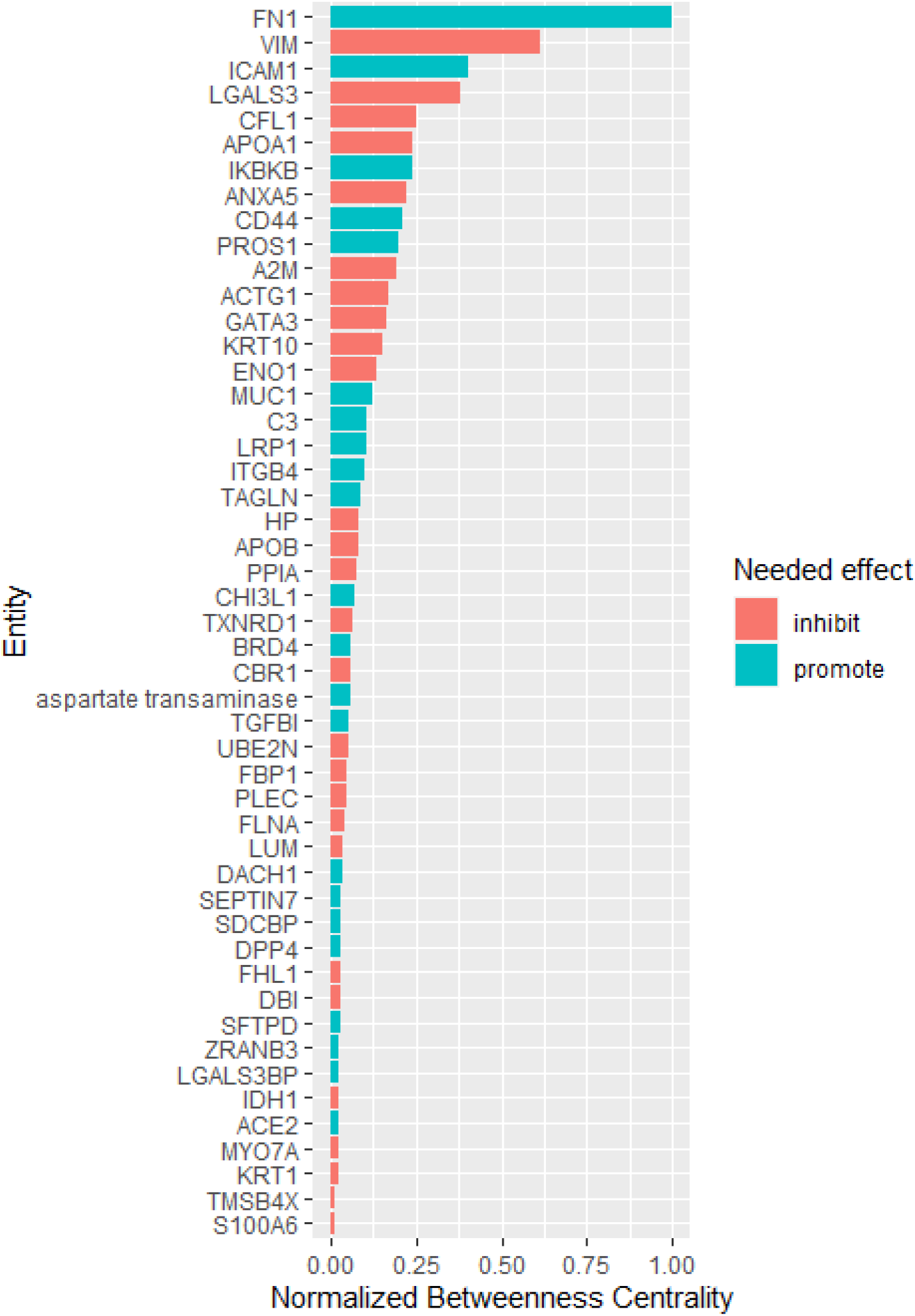
BALF network centrality nodes ranked by betweenness centrality. Betweenness centrality quantitatively describes how a node (in this case, a differentially expressed protein in the BALF proteome) mediates the interaction between communities of neighboring nodes in the network. Shown are 44 network entities with betweenness centrality >0.01, normalized to the maximum betweenness centrality present in the network. The betweenness centrality scores for all nodes were expressed as fractions of the maximum betweenness centrality present in the network. The (red and blue) colors indicate the needed effect (inhibition/induction) to restore these entities from COPD levels to the normal levels in healthy control subjects. The four nodes with ≥25% of the maximum betweenness centrality (fibronectin) with normalized betweenness centrality values representing a greater than the linear increase from the next lower ranking node are fibronectin, vimentin, intercellular adhesion molecule1 (ICAM1), and galectin-3. These potential key signaling mediators had a betweenness centrality of at least 25% of the maximum. Topological analysis of the interaction network regulatory interactions documented in the literature suggests that these proteins were central mediators of COPD.^57-61^ Colors indicate the needed effect to restore these entities to the normal levels in healthy control subjects. CANDO=computational analysis of novel drug opportunities COPD=chronic obstructive pulmonary disease

#### Network topological analysis

Four nodes representing proteins in the network stood out based on the normalized betweenness centrality values representing a greater than linear increase from the next lower ranking node: fibronectin, vimentin, intercellular adhesion molecule 1 (ICAM1), and galectin-3. These potentially key signaling mediators had a betweenness centrality of at least 25% of the maximum.

Analysis of the initial data reveals fibronectin and ICAM1 are reduced in COPD patients relative to healthy controls; thus, any candidate therapeutic should target an increase in their activity. The reverse is true for vimentin and galectin-3. We, therefore, sought drugs or combinations of drugs predicted to accomplish the appropriate activation or inhibition of the four most central nodes, specifically drugs that will lead to the promotion of central node proteins that were downregulated in the COPD cohort and inhibition of central node proteins that were overabundant in COPD. The idealized drug vector, therefore, constitutes interactions leading to desirable modulation of the central hub protein. CANDO identified 189 distinct drugs (**Figure 1, Table S7**S6) with relevance for COPD; 39 of these represented in the Elsevier Knowledge Graph ^31^ were analyzed for their enrichment for the desired agonist and antagonist effects on the most central entities in the protein regulatory network. Highly enriched drugs or drug pairs were are predicted to be more likely than randomly selected drugs to exert appropriate inhibition or promotion of the most central proteins. Two single drugs (fluocinolone acetonide and dexrazoxane) and 57 two-drug combinations were significantly enriched. Fluocinolone acetonide and dexrazoxane appeared in 54% and 46% of all significantly enriched 2-drug combinations, respectively, far greater than the other drugs appearing in these combinations (Figure 3). The combination of fluocinolone acetonide and dexrazoxane is the most enriched two-drug combination leading to an idealized drug vector.

**Figure 3.**
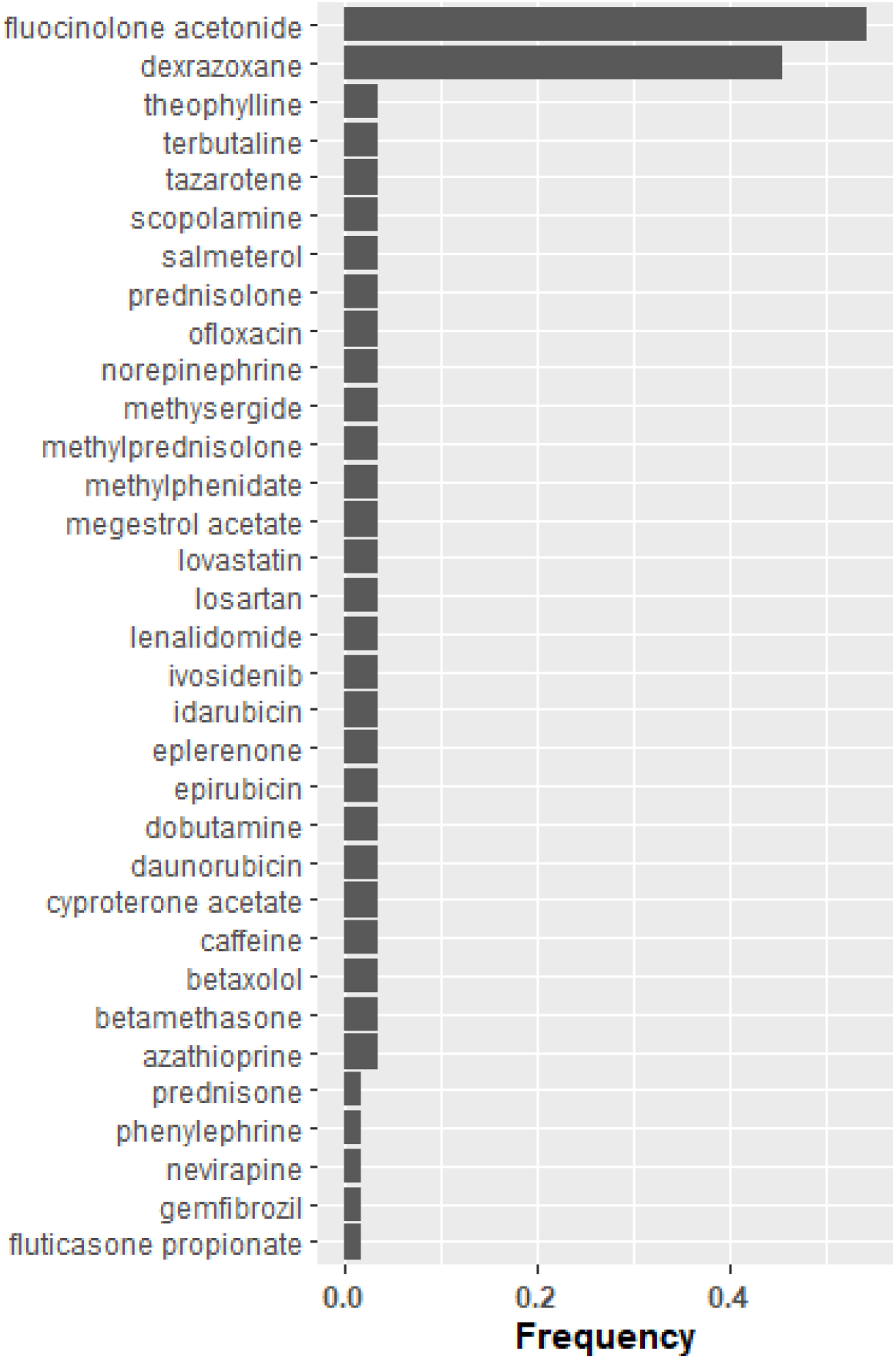
Drug frequency amongst idealized drug combinations predicted to modulate central proteins in COPD. The combinations of drugs initially identified by CANDO, which were predicted to activate or inhibit the four central nodes with the highest maximum betweenness centrality are listed (Figure 2). Drugs that lead to the promotion of central node proteins that were downregulated in the COPD cohort and inhibition of central node proteins (identified by network topological graph) that were overabundant in COPD. The idealized drug vector constitutes interactions leading to desirable modulation of the central hub protein. Representation of individual drug frequency among the 57 significantly enriched two- drug combinations (idealized drug vectors) out of the 39 proteins represented in the Elsevier Knowledge Graph are listed in descending order. Fluocinolone acetonide and dexrazoxane appeared in 54% and 46% of all significantly enriched two-drug combinations respectively, far greater than other drugs appearing in these combinations. The combination of fluocinolone acetonide and dexrazoxane is the most enriched two-drug combination leading to an idealized drug vector that most likely reverses the protein levels of the four central nodes to levels found in healthy control subjects. CANDO=computational analysis of novel drug opportunities COPD=chronic obstructive pulmonary disease

We additionally conducted a targeted query to assess the predicted effects of drugs commonly applied in pulmonary disease treatment on the most central proteins of this regulatory network (Figure 4 and Table S7). While some of these have been documented to have the desired effect on two of the central proteins, fibronectin or vimentin, all have been documented to have the opposite effect on at least one of the most central proteins. Therefore, they were not significantly enriched out of the set of all possible candidate drugs.

**Figure 4:**
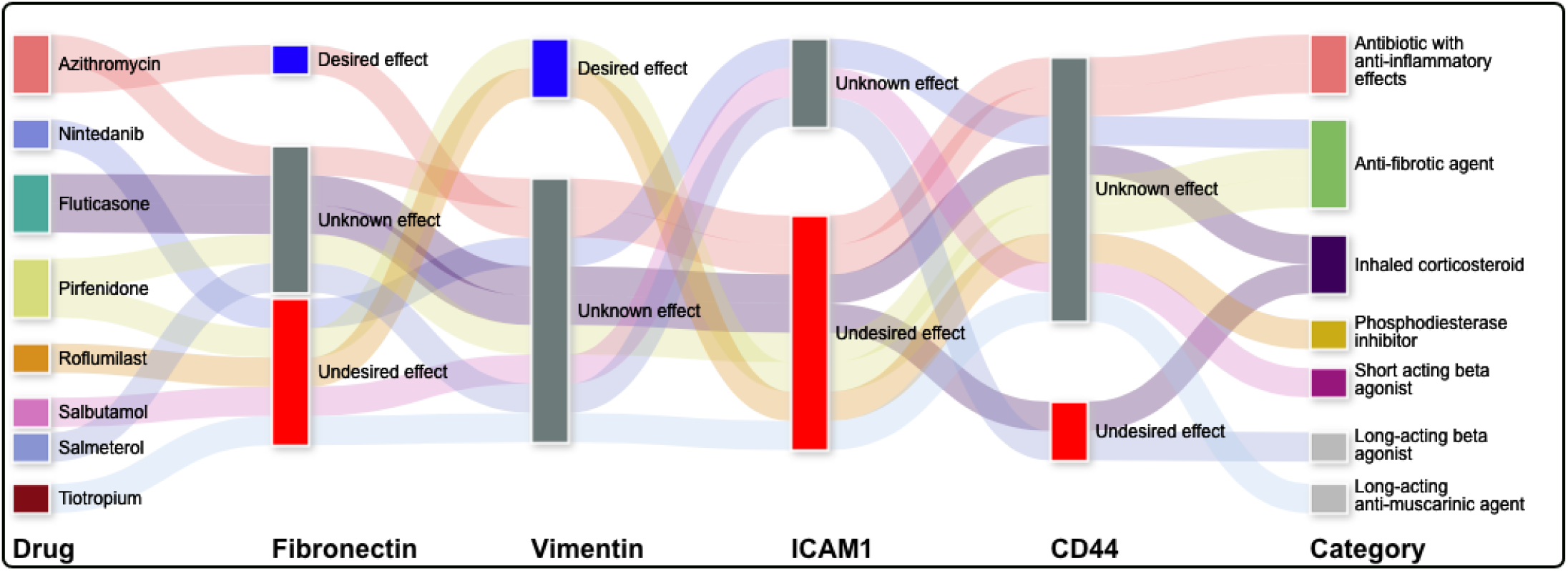
Commonly used pulmonary drugs and their putative effects on four central node proteins in COPD. A Sankey diagram categorizing the likely effects of putative drugs on four central node proteins in COPD is depicted. There are nine drugs on the left of the diagram used to treat different pulmonary diseases, with the corresponding drug classes displayed on the right side of the diagram. The effects of these drugs on four nodes (fibronectin, vimentin, intercellular adhesion molecule1 (ICAM1), and cd44) are detailed in the middle of the diagram, with broad lines connecting the proteins in the right to the putative effect (desired, unknown, undesired. While some of these have been documented to have the desired effect on fibronectin (promotion) or vimentin (inhibition), all have been reported to have the opposite effect on at least one of the most central proteins. This suggests using drugs commonly used to treat pulmonary disease, if repurposed for COPD, may have contrary effects on the mediators of the pathways involved in COPD, reinforcing the need to have a more nuanced approach to drug repurposing. CANDO=computational analysis of novel drug opportunities COPD=Chronic obstructive pulmonary disease

## Discussion

Our investigation of the COPD BALF proteome utilizing novel bioinformatic techniques revealed significant differences in proteins involved in multiple biological processes, including lung-specific mechanisms, protease/anti-protease homeostasis, immunoregulation, and the extracellular matrix. Proteomic profiling of the complex pathways implicated in COPD provides broader physiological exploration not provided by studying one entity at a time. We identified several differentially expressed proteins in COPD versus controls that, based on a review of published literature, have not been previously implicated in COPD etiology. This preliminary analysis illustrates how our BALF proteomic analysis represents a powerful approach to elucidate COPD pathogenesis and identify novel biomarkers.

Employing the bioinformatics tool DAVID and IPA, putative pathway networks were constructed based on the differentially expressed proteins in the BALF proteome that implicated multiple transcription factor pathways and disparate biological processes, such as extracellular space, proteolysis, extracellular matrix, cell adhesion, cytoskeleton, defense response, cell migration, and oxidation-reduction.

The CANDO platform identified 189 drug candidates that had similar protein interaction signatures based on the BALF proteome when compared to known drugs that are used to treat COPD. However, while most putative drug and protein interactions are likely inhibitors, the induction or inhibition of a target protein is indeterminable with solely the binding potential between drug and protein pairs.

Topological analysis of the interaction network connecting 233 proteins differentially expressed in COPD through regulatory interactions documented in the literature suggested that ICAM1^60^ and galectin-3^61^ are important central mediators of inflammation while both fibronectin^62,63^ and vimentin^64^ are central mediators of inflammation and fibrogenesis. This corroborates the results of the pathway enrichment analysis described above, and points to fibrosis and innate inflammation as important processes governing the pathogenesis and progression of COPD. A literature knowledge-based query (Elsevier Knowledge Graph) of drugs with desired drug-target interactions (generated using CANDO) identified putative drugs, such as anti-neoplastic, anti-fibrotic drugs, and regulators of inflammation, that would restore key central proteins to the levels characteristic of healthy controls. Our results also suggest currently utilized medications for COPD have disparate effects on the identified central node proteins that are key putative mediators of COPD pathogenesis and progression. In contrast, the corticosteroid fluocinolone acetonide^65^ and the cardioprotective agent dexrazoxane^66^ were highly enriched for the desired effects on central network entities, both individually and in combination. Fluocinolone acetonide is a stronger potentiator than other corticosteroids of the TGF-β pathway^67^ which is noted to be dysregulated in COPD^68^, and fluocinolone acetonide may be more effective than comparable corticosteroids in improved homeostasis in that pathway. Dexrazoxane^66^ is used to reduce cardiac toxicity associated with anthracycline-based chemotherapy agents by binding to iron and reducing reactive oxygen species; with oxidative stress as a significant factor in COPD pathogenesis^69^, antioxidative therapy may be beneficial.

The documented actions of these immunomodulators were predicted here to substantially counteract the observed dysregulation of centrally-connected proteins in COPD patients. The relatively high representation of immunomodulators among the candidate agents and the increased centrality of fibrosis-related proteins is consistent with the paradigm of airway remodeling as central to COPD pathology.^70^ With additional data, this regulatory circuit could be used as a testbed for computational evaluation of these and other candidate drug effects using network topological methods. ^71^

### Limitations and strengths

Our approach does has some limitations. The variability in how much BALF is recovered from each aliquot of saline infused in to the lower airway in COPD vs. control subjects are inherent in most BALF proteomic analyses. However, the BALF proteins were normalized to albumin BALF concentrations to account for the variability. The examination of protein levels without accounting for post-translational modifications, such as phosphorylation, may neglect important differences in protein interactions and activity, despite no significant differences in protein levels. Also, the BALF samples were from subjects in the COPD group who were ex-smokers. This exclusion limits the generalizability of our findings particularly current smokers, since the acute effects of tobacco smoke were excluded in our study design.

However, we confined our analysis to ex-smokers with moderate COPD to obtain some uniformity of the COPD phenotype and to avoid the acute inflammatory effects of current smoking. Future work on proteomic profiles will inform us of the difference between such profiles in current smokers and different stages of COPD.

### Comparison to previously published studies

A sputum proteomics study endeavored to identify COPD severity biomarkers by employing 2D gel electrophoresis and revealed 15 proteins that were significantly differentially expressed between healthy smoker controls and subjects with GOLD stage II; subsequently, 9 of the 15 candidate proteins were validated with Western Blot. Of the nine candidate proteins validated with Western Blot, seven were statistically significantly different between groups, specifically albumin, alpha-2-HS glycoprotein, transthyretin, PSP94, apolipoprotein A1, lipocalin-1, and PLUNC. ^72^ Employing quantitative ELISA data normalized for protein content, the investigators identified apolipoprotein A1 and lipocalin-1 as statistically differentially expressed in COPD. Although apolipoprotein A1 and lipocalin-1 were identified in our study of the BALF proteome, the proteins were not significantly differentially expressed, likely due to the differences in expression in the different biocompartments of sputum vs. bronchoalveolar lumen.

A 2D differential gel electrophoresis study and subsequent mass spectroscopy were performed by Ohlmeier et al., which compared healthy smokers, non-smokers, and smokers with GOLD stage II COPD and revealed a different set of 15 proteins that were differentially expressed between the groups.^73^ Of these proteins, polymeric immunoglobulin receptor levels in lung tissue and blood between the three groups were correlated with airflow obstruction.

In Lee et al., tumor-free lung tissue harvested from patients with lung cancer resection, when examined via 2D gel electrophoresis/MALDI-TOF-MS, revealed eight proteins that were upregulated in subjects with COPD compared to nonsmokers and ten significantly differentially expressed proteins between subjects with COPD and smoking subjects without COPD.^74^ Two of the identified proteins, matrix metalloprotease 13 (MMP13) and thioredoxin-like 2, were confirmed to be increased in COPD subjects with Western Blot and immunohistochemical staining, with MMP13 localized to the alveolar macrophage and type II pneumocytes and thioredoxin-like 2 found in the bronchial epithelium. Thioredoxin-like 2, which contains thioredoxin, was found in the BALF proteome but not significantly differentially expressed.

### However, MMP13 was not identified in our BALF study, likely due to differences in study populations and variable biocompartments

## Conclusion

In summary, our work provides a valuable pipepline for identifying many proteins associated with COPD pathogenesis that illustrate the complexity of the development of this disease, as well as identifying putative therapeutic treatment options using cutting-edge bioinformatics approaches. Identifying differentially expressed proteins will form the basis for future mechanistic studies of critical pathways and novel treatment discovery. Validation of our proposed therapeutic approach in animal models and pilot human studies are important next steps.

## Supporting information

Supplementary references

## Acknowledgments

The authors like to thank Catherine Wrona for her technical support.

