## Supplementary references for "Proteomic network analysis of bronchoalveolar lavage fluid in ex-smokers to discover implicated protein targets and novel drug treatments for chronic obstructive pulmonary disease"

Berenson CS, Garlipp MA, Grove LJ, Maloney J, Sethi S. Impaired phagocytosis of nontypeable Haemophilus influenzae by human alveolar macrophages in chronic obstructive pulmonary disease. *J Infect Dis.* 2006;194(10):1375-1384.

80.	Kaparianos A, Argyropoulou E. Local renin-angiotensin II systems, angiotensin-converting enzyme and its homologue ACE2: their potential role in the pathogenesis of chronic obstructive pulmonary diseases, pulmonary hypertension and acute respiratory distress syndrome. *Current Medicinal Chemistry.*18(23):3506-3515.

106.	Leikauf GD, Borchers MT, Prows DR, Simpson LG. Mucin apoprotein expression in COPD. *Chest.* 2002;121(5 Suppl):166S-182S.

107.	Freeman CM, Curtis JL, Chensue SW. CC chemokine receptor 5 and CXC chemokine receptor 6 expression by lung CD8+ cells correlates with chronic obstructive pulmonary disease severity. *American Journal of Pathology.* 2007;171(3):767-776.

108.	Su YW, Xu YJ, Liu XS. Quantitative differentiation of dendritic cells in lung tissues of smokers with and without chronic obstructive pulmonary disease. *Chinese Medical Journal.* 2010;123(12):1500-1504.

109.	Tsoumakidou M, Koutsopoulos AV, Tzanakis N, et al. Decreased small airway and alveolar CD83+ dendritic cells in COPD. *Chest.* 2009;136(3):726-733.

110.	Verhoeven GT, Hegmans JP, Mulder PG, Bogaard JM, Hoogsteden HC, Prins JB. Effects of fluticasone propionate in COPD patients with bronchial hyperresponsiveness. *Thorax.* 2002;57(8):694-700.
