## Supplementary material for "Proteomic network analysis of bronchoalveolar lavage fluid in ex-smokers to discover implicated protein targets and novel drug treatments for chronic obstructive pulmonary disease": Figure S5

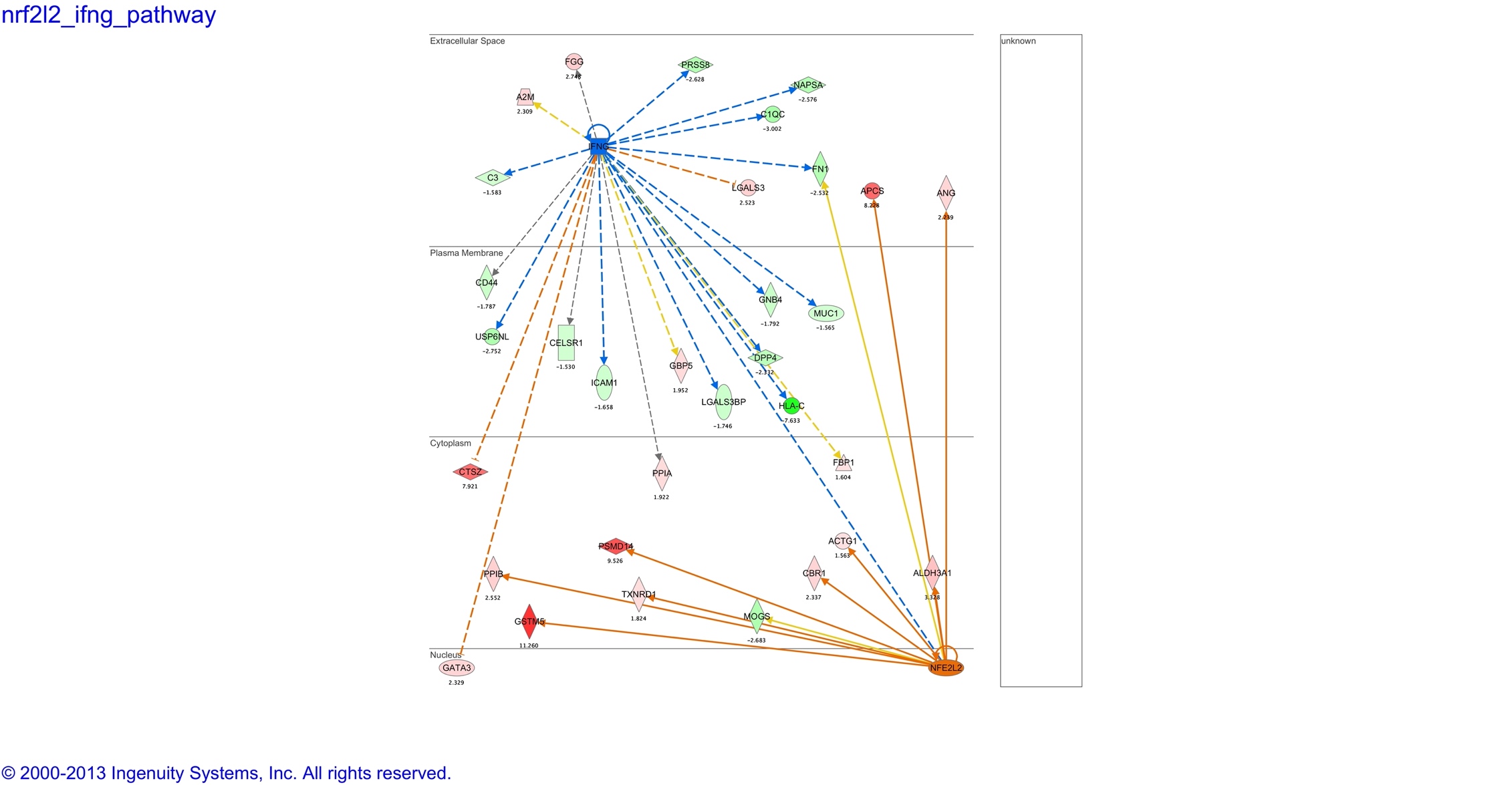

- **Figure S5:**
- **IPA network putative upstream regulators**

Putative upstream regulators of the proteins that were significantly differentially expressed between the cohorts were associated with a predicted downregulation and upregulation of interferon gamma and nuclear factor erythroid 2-related 2 (NRF2) respectively.

Solid lines indicated a direct interaction between proteins, while dotted lines indicate an indirect association between two proteins. Proteins upregulated in the BALF dataset and are predicted are shaded in red and proteins downregulated in the BALF dataset are shaded in green. Interferon gamma is shaded in blue to denote a putative downregulation of the protein while NRF2 is shaded in orange to denote a putative upregulation of the protein based on the IPA relational database. The darker shading to lighter shading corresponds to decreasing expression intensity. The lines colored with orange shading correspond to interactions of the upstream regulator leading to increased protein synthesis of the downstream protein. The lines colored with blue shading correspond to interactions of the upstream regulator leading to decreased protein synthesis of the downstream protein. Lines colored with yellow shading indicate a downstream protein expression level that is discordant with the putative interaction from the upstream protein.

IPA=Ingenuity Pathway Analysis, BALF= Bronchoalveolar Lavage Fluid]
