## Supplementary material for "Proteomic network analysis of bronchoalveolar lavage fluid in ex-smokers to discover implicated protein targets and novel drug treatments for chronic obstructive pulmonary disease": Figure S4

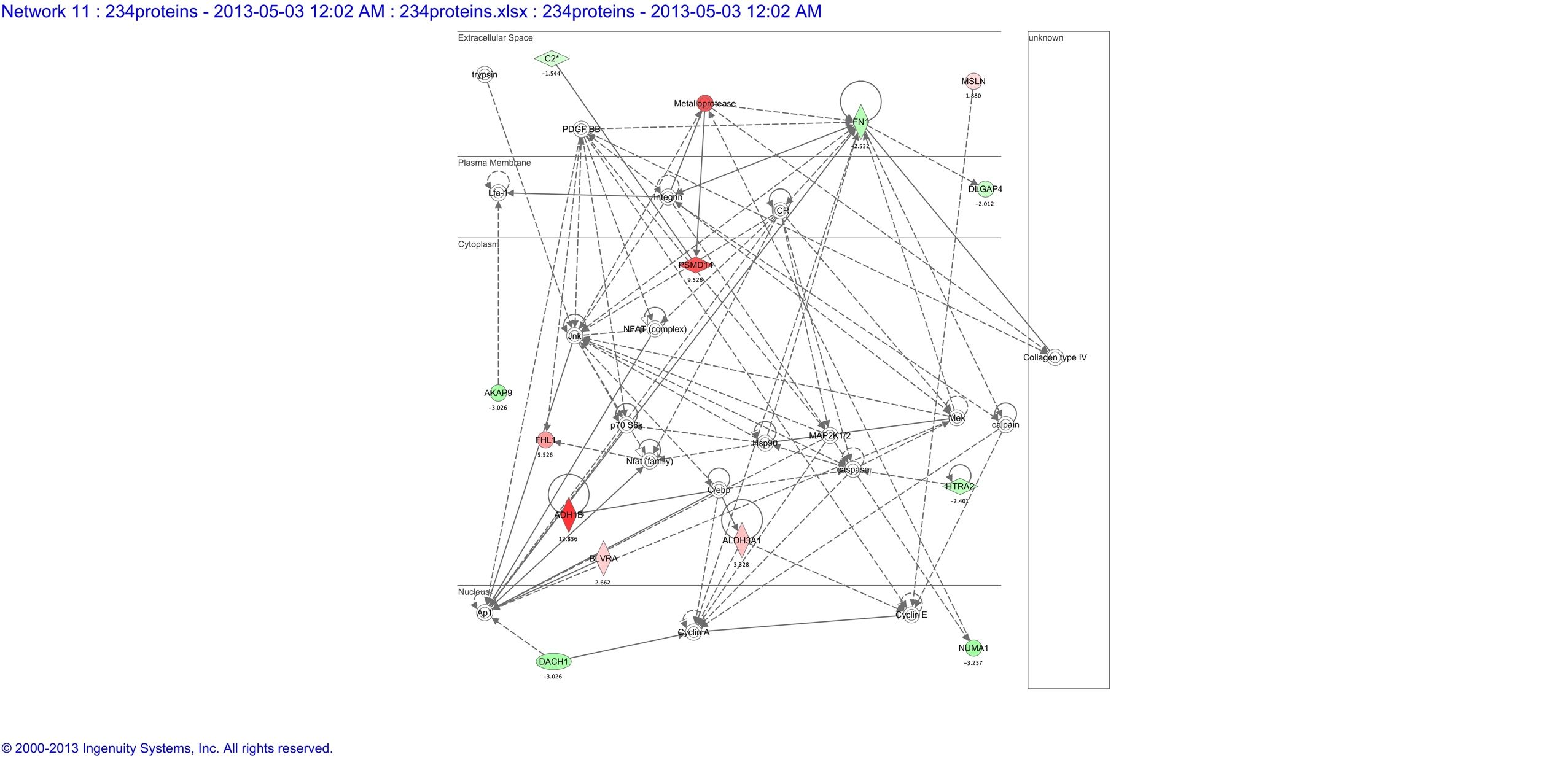

**Figure S4:**

**IPA network 11: cell cycle, visual system development and function, hair and skin development and function**

Functional annotation networks from IPA that show relationships among the genes that in IPA’s relational database are related to cell cycle, visual system development and function, hair and skin development and function.

Solid lines indicated a direct interaction between proteins, while dotted lines indicate an indirect association between two proteins. Proteins upregulated in the BALF dataset are shaded in red and proteins downregulated in the BALF dataset are shaded in green. The darker shading to lighter shading corresponds to decreasing expression intensity.

IPA=Ingenuity Pathway Analysis, BALF= Bronchoalveolar Lavage Fluid
