## Supplementary material for "Proteomic network analysis of bronchoalveolar lavage fluid in ex-smokers to discover implicated protein targets and novel drug treatments for chronic obstructive pulmonary disease": Figure S3

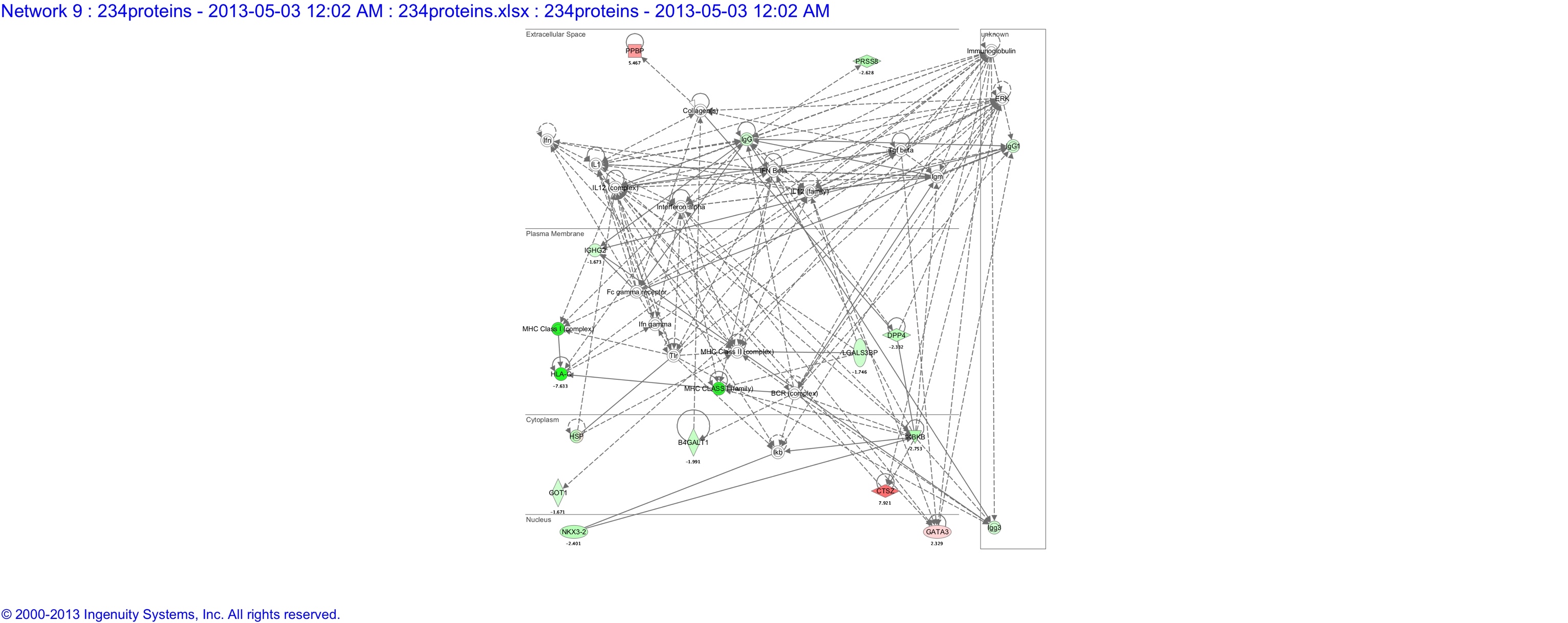

**Figure S3:**

**IPA network 9: cellular movement, hematological system development and function, immune cell trafficking**

Functional annotation networks from IPA that show relationships among the genes that in IPA’s relational database are related to cellular movement, hematological system development and function, and immune cell trafficking.

Solid lines indicated a direct interaction between proteins, while dotted lines indicate an indirect association between two proteins. Proteins upregulated in the BALF dataset are shaded in red and proteins downregulated in the BALF dataset are shaded in green. The darker shading to lighter shading corresponds to decreasing expression intensity.

IPA=Ingenuity Pathway Analysis, BALF= Bronchoalveolar Lavage Fluid
