## Supplementary material for "Proteomic network analysis of bronchoalveolar lavage fluid in ex-smokers to discover implicated protein targets and novel drug treatments for chronic obstructive pulmonary disease": Figure S2

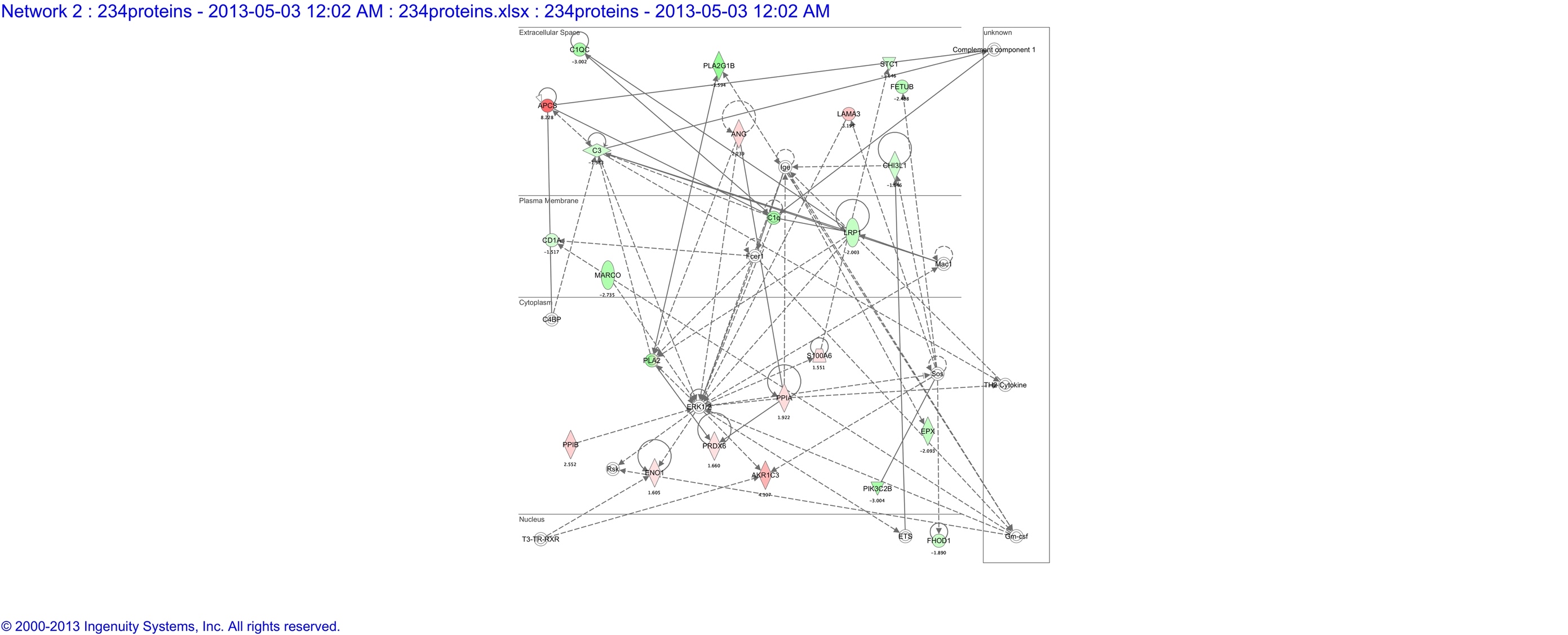

**Figure S2:**

**IPA network 2: cell death and survival, drug metabolism, small molecule biochemistry**

Functional annotation networks from IPA that show relationships among the genes that in IPA’s relational database are related to cell death and survival, drug metabolism and small molecule biochemistry.

Solid lines indicated a direct interaction between proteins, while dotted lines indicate an indirect association between two proteins. Proteins upregulated in the BALF dataset are shaded in red and proteins downregulated in the BALF dataset are shaded in green. The darker shading to lighter shading corresponds to decreasing expression intensity.

IPA=Ingenuity Pathway Analysis, BALF= Bronchoalveolar Lavage Fluid
