## Supplementary material for "Proteomic network analysis of bronchoalveolar lavage fluid in ex-smokers to discover implicated protein targets and novel drug treatments for chronic obstructive pulmonary disease": Figure S1

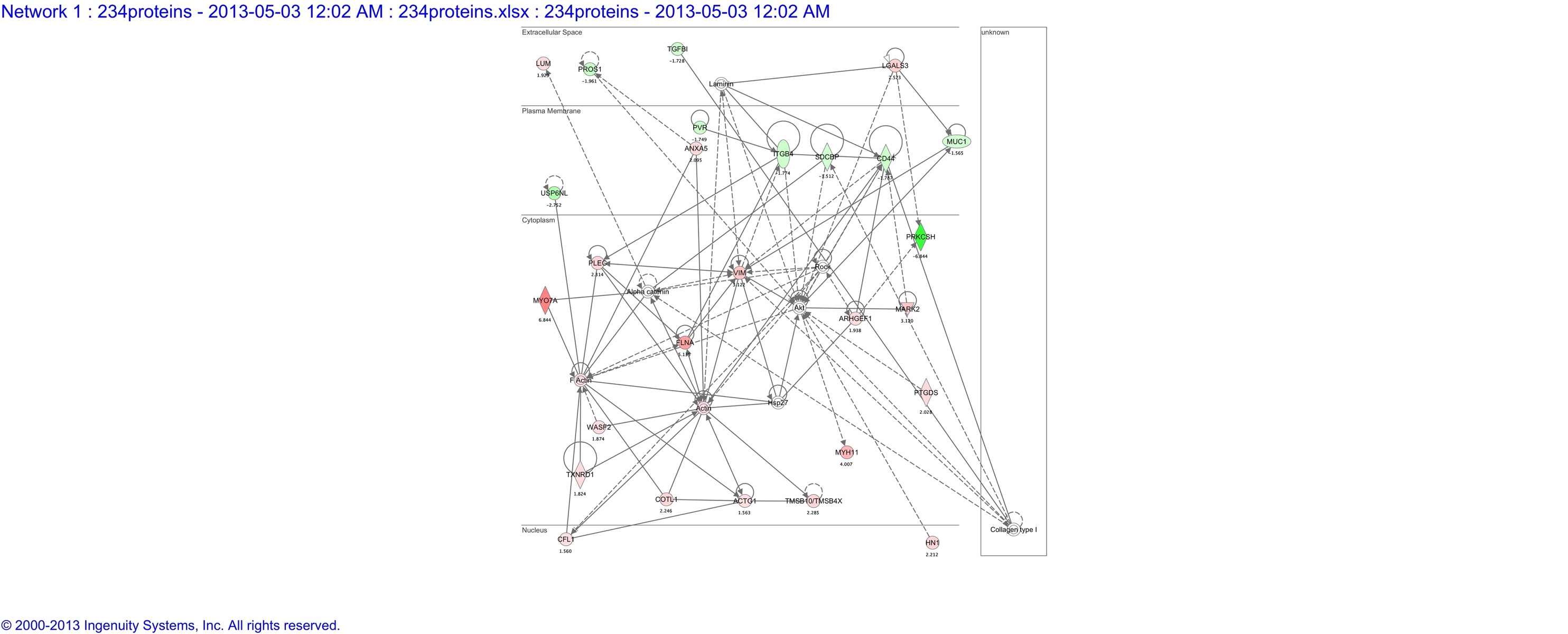

**Figure S1:**

**IPA Network 1: Cellular Movement, Inflammatory Response, Cardiovascular System Development and Function**

Functional annotation networks from IPA show relationships among the genes that in IPA’s relational database are related to cellular movement, inflammatory response and cardiovascular system development and function.

Solid lines indicated a direct interaction between proteins, while dotted lines indicate an indirect association between two proteins. Proteins upregulated in the BALF dataset are shaded in red and proteins downregulated in the BALF dataset are shaded in green. The darker shading to lighter shading corresponds to decreasing expression intensity.

IPA=Ingenuity Pathway Analysis, BALF= Bronchoalveolar Lavage Fluid
