## Supplementary material for "Proteomic network analysis of bronchoalveolar lavage fluid in ex-smokers to discover implicated protein targets and novel drug treatments for chronic obstructive pulmonary disease": Table S8

**Predicted interactions of drugs treating respiratory diseases and central node proteins**

Representative drugs treating respiratory disease from selected categories showing their predicted interactions with the most central nodes entities of Figure 4

| - **Drug** | - **Category** | - **Undesired effect** | - **Desired effect** |
| --- | --- | --- | --- |
| - azithromycin | - antibiotic with anti-inflammatory effects | - VIM, ICAM1 | - FN1 |
| - BIBF 1120 | - anti-fibrotic agent | - FN1 | - -- |
| - fluticasone | - inhaled corticosteroid | - ICAM1, CD44 | - -- |
| - pirfenidone | - anti-fibrotic agent | - FN1, ICAM1 | - VIM |
| - roflumilast | - phosphodiesterase inhibitor | - ICAM1, FN1 | - VIM |
| - salbutamol | - short acting beta agonist | - FN1 | - -- |
| - salmeterol | - long-acting beta agonist | - CD44 | - -- |
| - tiotropium | - long-acting anti-muscarinic agent | - FN1, ICAM1 | - -- |
