## Supplementary material for "Proteomic network analysis of bronchoalveolar lavage fluid in ex-smokers to discover implicated protein targets and novel drug treatments for chronic obstructive pulmonary disease": Table S7

**Computational drug prediction CANDO**

- (score1= refers to the consensus score or number of times the compound shows up in the top 30 most similar drugs used to treat COPD

score2= the average of the ranks for 'score1’

probability= the binomial distribution derived probability of achieving ‘score1’ by chance based on the number of drugs associated with COPD, the total number of drugs in the library, and the number of most similar drugs to consider (in this case, 30).

name= generic name of the candidate drug)

CANDO= Computational Analysis of Novel Drug Opportunities

COPD= chronic obstructive pulmonary disease

| **Rank** | **Score1 (consensus score)** | **Score2 (average ranks score)** | **Probability** | **Name** |
| --- | --- | --- | --- | --- |
| 1 | 12 | 10.1 | 1.11E-16 | clobetasol_propionate |
| 2 | 12 | 11.8 | 1.11E-16 | clobetasol |
| 3 | 12 | 12.7 | 1.11E-16 | rimexolone |
| 4 | 11 | 12.5 | 4.88E-15 | deflazacort |
| 5 | 10 | 7.2 | 2.07E-13 | loteprednol_etabonate |
| 6 | 10 | 10.4 | 2.07E-13 | desoximetasone |
| 7 | 10 | 13.1 | 2.07E-13 | loteprednol |
| 8 | 10 | 13.4 | 2.07E-13 | meprednisone |
| 9 | 10 | 14.4 | 2.07E-13 | amcinonide |
| 10 | 9 | 13.2 | 7.66E-12 | fluclorolone_acetonide |
| 11 | 9 | 15.1 | 7.66E-12 | fluorometholone |
| 12 | 9 | 17.7 | 7.66E-12 | clobetasone |
| 13 | 8 | 14.2 | 2.48E-10 | ulobetasol |
| 14 | 8 | 16.1 | 2.48E-10 | procaterol |
| 15 | 8 | 18.1 | 2.48E-10 | desonide |
| 16 | 7 | 11 | 6.95E-09 | prednicarbate |
| 17 | 7 | 11.6 | 6.95E-09 | tezacaftor |
| 18 | 7 | 16 | 6.95E-09 | cyproterone_acetate |
| 19 | 7 | 24.3 | 6.95E-09 | drometrizole_trisiloxane |
| 20 | 6 | 14.2 | 1.67E-07 | nadolol |
| 21 | 6 | 14.5 | 1.67E-07 | hydrocortamate |
| 22 | 6 | 19.7 | 1.67E-07 | hydrocortisone_butyrate |
| 23 | 6 | 23.2 | 1.67E-07 | flurandrenolide |
| 24 | 5 | 3 | 3.39E-06 | isoprenaline |
| 25 | 5 | 4.6 | 3.39E-06 | epinephrine |
| 26 | 5 | 5.6 | 3.39E-06 | orciprenaline |
| 27 | 5 | 6.6 | 3.39E-06 | isoetharine |
| 28 | 5 | 7.6 | 3.39E-06 | carbidopa |
| 29 | 5 | 9.2 | 3.39E-06 | gemfibrozil |
| 30 | 5 | 9.8 | 3.39E-06 | phenylephrine |
| 31 | 5 | 9.8 | 3.39E-06 | methyldopa |
| 32 | 5 | 10.8 | 3.39E-06 | carteolol |
| 33 | 5 | 11.2 | 3.39E-06 | paramethasone_acetate |
| 34 | 5 | 11.8 | 3.39E-06 | clocortolone |
| 35 | 5 | 12.2 | 3.39E-06 | arbutamine |
| 36 | 5 | 12.4 | 3.39E-06 | pindolol |
| 37 | 5 | 13.4 | 3.39E-06 | hydrocortisone_acetate |
| 38 | 5 | 13.8 | 3.39E-06 | levobunolol |
| 39 | 5 | 14.6 | 3.39E-06 | difluocortolone |
| 40 | 5 | 17.6 | 3.39E-06 | propofol |
| 41 | 5 | 18.6 | 3.39E-06 | celiprolol |
| 42 | 5 | 18.8 | 3.39E-06 | levonordefrin |
| 43 | 5 | 20.8 | 3.39E-06 | tapentadol |
| 44 | 5 | 20.8 | 3.39E-06 | segesterone_acetate |
| 45 | 4 | 9.2 | 5.72E-05 | betamethasone |
| 46 | 4 | 9.2 | 5.72E-05 | methylprednisolone_aceponate |
| 47 | 4 | 10.2 | 5.72E-05 | dexamethasone |
| 48 | 4 | 11 | 5.72E-05 | naldemedine |
| 49 | 4 | 11.2 | 5.72E-05 | elvitegravir |
| 50 | 4 | 14 | 5.72E-05 | difluprednate |
| 51 | 4 | 15.8 | 5.72E-05 | deferiprone |
| 52 | 4 | 17 | 5.72E-05 | methylprednisolone |
| 53 | 4 | 17.5 | 5.72E-05 | prednisolone |
| 54 | 4 | 18 | 5.72E-05 | tamsulosin |
| 55 | 4 | 20.5 | 5.72E-05 | fluprednisolone |
| 56 | 4 | 22.2 | 5.72E-05 | canrenoic_acid |
| 57 | 4 | 22.5 | 5.72E-05 | etidocaine |
| 58 | 4 | 23 | 5.72E-05 | mephenesin |
| 59 | 4 | 24.5 | 5.72E-05 | hydrocortisone_valerate |
| 60 | 4 | 25.5 | 5.72E-05 | halcinonide |
| 61 | 4 | 26 | 5.72E-05 | desvenlafaxine |
| 62 | 3 | 5 | 7.80E-04 | fexofenadine |
| 63 | 3 | 7 | 7.80E-04 | pioglitazone |
| 64 | 3 | 7.7 | 7.80E-04 | laropiprant |
| 65 | 3 | 8.3 | 7.80E-04 | terfenadine |
| 66 | 3 | 8.7 | 7.80E-04 | cyclandelate |
| 67 | 3 | 9.7 | 7.80E-04 | dobutamine |
| 68 | 3 | 10 | 7.80E-04 | mometasone_furoate |
| 69 | 3 | 10.3 | 7.80E-04 | fentanyl |
| 70 | 3 | 11 | 7.80E-04 | amiodarone |
| 71 | 3 | 11.3 | 7.80E-04 | siponimod |
| 72 | 3 | 12.3 | 7.80E-04 | homatropine_methylbromide |
| 73 | 3 | 14 | 7.80E-04 | metaraminol |
| 74 | 3 | 14 | 7.80E-04 | flunisolide |
| 75 | 3 | 14.3 | 7.80E-04 | meradimate |
| 76 | 3 | 14.7 | 7.80E-04 | fluocinonide |
| 77 | 3 | 15 | 7.80E-04 | masoprocol |
| 78 | 3 | 15 | 7.80E-04 | fluocinolone_acetonide |
| 79 | 3 | 15.3 | 7.80E-04 | loperamide |
| 80 | 3 | 16 | 7.80E-04 | hydrocortisone_cypionate |
| 81 | 3 | 16 | 7.80E-04 | piritramide |
| 82 | 3 | 16.3 | 7.80E-04 | darifenacin |
| 83 | 3 | 17 | 7.80E-04 | ebastine |
| 84 | 3 | 18 | 7.80E-04 | guaifenesin |
| 85 | 3 | 18.7 | 7.80E-04 | zolmitriptan |
| 86 | 3 | 19 | 7.80E-04 | trospium |
| 87 | 3 | 19.3 | 7.80E-04 | darolutamide |
| 88 | 3 | 19.7 | 7.80E-04 | olmesartan |
| 89 | 3 | 20.3 | 7.80E-04 | megestrol_acetate |
| 90 | 3 | 20.7 | 7.80E-04 | levocabastine |
| 91 | 3 | 23 | 7.80E-04 | benserazide |
| 92 | 3 | 26 | 7.80E-04 | stiripentol |
| 93 | 3 | 27 | 7.80E-04 | dipivefrin |
| 94 | 3 | 29 | 7.80E-04 | norepinephrine |
| 95 | 2 | 1 | 8.29E-03 | oxtriphylline |
| 96 | 2 | 2 | 8.29E-03 | bromotheophylline |
| 97 | 2 | 2 | 8.29E-03 | methscopolamine_bromide |
| 98 | 2 | 2 | 8.29E-03 | butylscopolamine |
| 99 | 2 | 2.5 | 8.29E-03 | diethylamino_hydroxybenzoyl_hexyl_benzoate |
| 100 | 2 | 3 | 8.29E-03 | caffeine |
| 101 | 2 | 3 | 8.29E-03 | methscopolamine |
| 102 | 2 | 4 | 8.29E-03 | xanthinol |
| 103 | 2 | 4 | 8.29E-03 | dopexamine |
| 104 | 2 | 4.5 | 8.29E-03 | difenoxin |
| 105 | 2 | 4.5 | 8.29E-03 | scopolamine |
| 106 | 2 | 5 | 8.29E-03 | enprofylline |
| 107 | 2 | 5 | 8.29E-03 | oxyphenonium |
| 108 | 2 | 5.5 | 8.29E-03 | cortisone_acetate |
| 109 | 2 | 5.5 | 8.29E-03 | labetalol |
| 110 | 2 | 6 | 8.29E-03 | pentoxifylline |
| 111 | 2 | 6 | 8.29E-03 | oxybutynin |
| 112 | 2 | 7 | 8.29E-03 | dyphylline |
| 113 | 2 | 7 | 8.29E-03 | methylphenidate |
| 114 | 2 | 8 | 8.29E-03 | temozolomide |
| 115 | 2 | 8 | 8.29E-03 | cyclopentolate |
| 116 | 2 | 8 | 8.29E-03 | dexmethylphenidate |
| 117 | 2 | 9 | 8.29E-03 | enoxacin |
| 118 | 2 | 9 | 8.29E-03 | lemborexant |
| 119 | 2 | 9 | 8.29E-03 | diflorasone |
| 120 | 2 | 9 | 8.29E-03 | fluocortolone |
| 121 | 2 | 9.5 | 8.29E-03 | apremilast |
| 122 | 2 | 9.5 | 8.29E-03 | mepenzolate |
| 123 | 2 | 9.5 | 8.29E-03 | mebeverine |
| 124 | 2 | 9.5 | 8.29E-03 | etofamide |
| 125 | 2 | 10 | 8.29E-03 | tipiracil |
| 126 | 2 | 10 | 8.29E-03 | flumethasone |
| 127 | 2 | 10.5 | 8.29E-03 | diphenoxylate |
| 128 | 2 | 10.5 | 8.29E-03 | ambenonium |
| 129 | 2 | 12 | 8.29E-03 | dexrazoxane |
| 130 | 2 | 12 | 8.29E-03 | rosiglitazone |
| 131 | 2 | 12.5 | 8.29E-03 | halofantrine |
| 132 | 2 | 13 | 8.29E-03 | daunorubicin |
| 133 | 2 | 13 | 8.29E-03 | dicloxacillin |
| 134 | 2 | 13.5 | 8.29E-03 | permethrin |
| 135 | 2 | 13.5 | 8.29E-03 | trimethaphan |
| 136 | 2 | 14 | 8.29E-03 | tinidazole |
| 137 | 2 | 14 | 8.29E-03 | penbutolol |
| 138 | 2 | 14 | 8.29E-03 | oxyphencyclimine |
| 139 | 2 | 15 | 8.29E-03 | acetazolamide |
| 140 | 2 | 15 | 8.29E-03 | methylergometrine |
| 141 | 2 | 15 | 8.29E-03 | cloxacillin |
| 142 | 2 | 15 | 8.29E-03 | ecamsule |
| 143 | 2 | 15 | 8.29E-03 | sonidegib |
| 144 | 2 | 15.5 | 8.29E-03 | cefpirome |
| 145 | 2 | 15.5 | 8.29E-03 | zofenopril |
| 146 | 2 | 15.5 | 8.29E-03 | panobinostat |
| 147 | 2 | 16 | 8.29E-03 | methimazole |
| 148 | 2 | 16.5 | 8.29E-03 | troglitazone |
| 149 | 2 | 16.5 | 8.29E-03 | flucloxacillin |
| 150 | 2 | 17 | 8.29E-03 | levofloxacin |
| 151 | 2 | 17 | 8.29E-03 | sumatriptan |
| 152 | 2 | 17 | 8.29E-03 | pentoxyverine |
| 153 | 2 | 17.5 | 8.29E-03 | elagolix |
| 154 | 2 | 17.5 | 8.29E-03 | hexylcaine |
| 155 | 2 | 18 | 8.29E-03 | ofloxacin |
| 156 | 2 | 18 | 8.29E-03 | alclometasone |
| 157 | 2 | 19 | 8.29E-03 | lomefloxacin |
| 158 | 2 | 19 | 8.29E-03 | ioflupane_i-123 |
| 159 | 2 | 19 | 8.29E-03 | losartan |
| 160 | 2 | 19.5 | 8.29E-03 | bemotrizinol |
| 161 | 2 | 20 | 8.29E-03 | epirubicin |
| 162 | 2 | 20 | 8.29E-03 | benzethonium |
| 163 | 2 | 20 | 8.29E-03 | benazepril |
| 164 | 2 | 20 | 8.29E-03 | mepyramine |
| 165 | 2 | 20.5 | 8.29E-03 | maraviroc |
| 166 | 2 | 20.5 | 8.29E-03 | methysergide |
| 167 | 2 | 20.5 | 8.29E-03 | dextropropoxyphene |
| 168 | 2 | 20.5 | 8.29E-03 | ritodrine |
| 169 | 2 | 21 | 8.29E-03 | doxorubicin |
| 170 | 2 | 21 | 8.29E-03 | cinchocaine |
| 171 | 2 | 21 | 8.29E-03 | cefapirin |
| 172 | 2 | 21.5 | 8.29E-03 | tegaserod |
| 173 | 2 | 21.5 | 8.29E-03 | nomegestrol |
| 174 | 2 | 22 | 8.29E-03 | dacarbazine |
| 175 | 2 | 22.5 | 8.29E-03 | oxeladin |
| 176 | 2 | 23 | 8.29E-03 | deutetrabenazine |
| 177 | 2 | 23 | 8.29E-03 | tropicamide |
| 178 | 2 | 23 | 8.29E-03 | triamcinolone |
| 179 | 2 | 23.5 | 8.29E-03 | lovastatin |
| 180 | 2 | 24 | 8.29E-03 | methazolamide |
| 181 | 2 | 25 | 8.29E-03 | pefloxacin |
| 182 | 2 | 26 | 8.29E-03 | nalidixic_acid |
| 183 | 2 | 26 | 8.29E-03 | norgestimate |
| 184 | 2 | 26.5 | 8.29E-03 | gestrinone |
| 185 | 2 | 26.5 | 8.29E-03 | ergometrine |
| 186 | 2 | 27 | 8.29E-03 | dorzolamide |
| 187 | 2 | 27 | 8.29E-03 | ethylhexyl_methoxycrylene |
| 188 | 2 | 28 | 8.29E-03 | lenalidomide |
| 189 | 2 | 29 | 8.29E-03 | idarubicin |
