## Supplementary material for "Proteomic network analysis of bronchoalveolar lavage fluid in ex-smokers to discover implicated protein targets and novel drug treatments for chronic obstructive pulmonary disease": Table S6

- Table S5:
- **Transcription factors associated with binding sites on genes from differentially expressed proteins in BALF.** Transcription factors that have association with binding sites on genes from differentially expressed proteins in BALF as noted by DAVID [ref].

| Transcription factor with binding sites in the genes represented in BALF | Number of the differentially expressed proteins that have corresponding gene binding sites with transcription factor | % of dataset in DAVID database | p-value  (Fisher exact) |
| --- | --- | --- | --- |
| AREB6 | 166 | 79.43 | 0.009 |
| SRF | 148 | 70.81 | 0.000 |
| AML1 | 147 | 70.33 | 0.089 |
| P53 | 127 | 60.77 | 0.061 |
| AP4 | 125 | 59.81 | 0.053 |
| LMO2COM | 122 | 58.37 | 0.004 |
| SREBP1 | 118 | 56.46 | 0.046 |
| PAX2 | 113 | 54.07 | 0.073 |
| USF | 113 | 54.07 | 0.081 |
| PAX5 | 112 | 53.59 | 0.034 |
| GCNF | 112 | 53.59 | 0.049 |
| STAT5A | 111 | 53.11 | 0.074 |
| MRF2 | 110 | 52.63 | 0.001 |
| HTF | 110 | 52.63 | 0.010 |
| TAXCREB | 109 | 52.15 | 0.021 |
| CEBPB | 109 | 52.15 | 0.041 |
| AHRARNT | 105 | 50.24 | 0.031 |
| FOXO4 | 105 | 50.24 | 0.073 |
| RP58 | 104 | 49.76 | 0.032 |
| STAT3 | 101 | 48.33 | 0.004 |
| BACH1 | 101 | 48.33 | 0.028 |
| HNF4 | 100 | 47.85 | 0.010 |
| GFI1 | 97 | 46.41 | 0.002 |
| NFKB | 97 | 46.41 | 0.071 |
| STAT1 | 96 | 45.93 | 0.000 |
| FREAC3 | 95 | 45.45 | 0.001 |
| CREBP1 | 95 | 45.45 | 0.033 |
| OCT | 94 | 44.98 | 0.025 |
| CDPCR3HD | 92 | 44.02 | 0.019 |
| HAND1E47 | 92 | 44.02 | 0.052 |
| RSRFC4 | 91 | 43.54 | 0.091 |
| SOX9 | 90 | 43.06 | 0.028 |
| HFH1 | 90 | 43.06 | 0.044 |
| CMYB | 89 | 42.58 | 0.014 |
| GATA | 89 | 42.58 | 0.050 |
| HSF2 | 88 | 42.11 | 0.012 |
| IK3 | 88 | 42.11 | 0.058 |
| HOX13 | 88 | 42.11 | 0.098 |
| RORA1 | 87 | 41.63 | 0.087 |
| HLF | 86 | 41.15 | 0.009 |
| FOXO1 | 85 | 40.67 | 0.026 |
| TGIF | 84 | 40.19 | 0.060 |
| POU6F1 | 84 | 40.19 | 0.068 |
| E4BP4 | 84 | 40.19 | 0.100 |
| MIF1 | 83 | 39.71 | 0.074 |
| STAT | 82 | 39.23 | 0.001 |
| CP2 | 82 | 39.23 | 0.020 |
| MSX1 | 81 | 38.76 | 0.098 |
| LYF1 | 80 | 38.28 | 0.032 |
| HNF3B | 80 | 38.28 | 0.091 |
| NFKAPPAB | 79 | 37.80 | 0.004 |
| NFE2 | 70 | 33.49 | 0.094 |
| FOXD3 | 69 | 33.01 | 0.061 |
| IK2 | 62 | 29.67 | 0.054 |
| TAL1BETAE47 | 61 | 29.19 | 0.041 |
| HSF1 | 61 | 29.19 | 0.087 |
| ZIC2 | 53 | 25.36 | 0.016 |
| GATA3 | 51 | 24.40 | 0.034 |
| MAX | 37 | 17.70 | 0.089 |

- **Table S6: Top functional networks of differentially expressed molecules in the BALF proteome.** The top biological functions associated with molecular pathways imputed with IPA that are significantly associated differentially expressed molecules measured in the BALF proteome. Red represents upregulated, and green represents downregulated proteins. The networks are collections of interconnected molecules assembled by a network algorithm. Each connection represents known relationships between the molecules, found in the Ingenuity Knowledge Base. The score is the degree of relevance of network eligible molecules to the BALF dataset.  The score takes into account the number of network eligible molecules in the network and its size, as well as the total number of network eligible molecules analyzed and the total number of molecules in the Ingenuity Knowledge Base that could potentially be included in networks.  The network score is based on the hypergeometric distribution and is calculated with the right-tailed Fisher's Exact Test: Score=-log(Fisher's Exact test result). Focus Molecules are the number of proteins identified in the BALF proteome that is found in the network.

| **ID** | Molecules in network | Score | Focus molecules | Top biological functions associated with the molecular network |
| --- | --- | --- | --- | --- |
| **1** | ACTG1, Actin, Akt, Alpha catenin, ANXA5, ARHGEF1, CD44, CFL1, Collagen type I, COTL1, F Actin, FLNA, HN1, Hsp27, ITGB4, Laminin, LGALS3, LUM, MARK2, MUC1, MYH11, MYO7A, PLEC, PRKCSH, PROS1, PTGDS, PVR, Rock, SDCBP, TGFBI, TMSB10/TMSB4X, TXNRD1, USP6NL, VIM, WASF2 | 51 | 27 | Cellular Movement, Inflammatory Response, Cardiovascular System Development and Function |
| **2** | AKR1C3, ANG, APCS, C3, C1q, C1QC, C4BP, CD1A, CHI3L1, Complement component 1, ENO1, EPX, ERK1/2, ETS, Fcer1, FETUB, FHOD1, Gm-csf, Ige, LAMA3, LRP1, Mac1, MARCO, PIK3C2B, PLA2, PLA2G1B, PPIA, PPIB, PRDX6, Rsk, S100A6, Sos, STC1, T3-TR-RXR, TH2 Cytokine | 36 | 21 | Cell Death and Survival, Drug Metabolism, Small Molecule Biochemistry |
| **3** | APC, BOD1L1, C10orf116, C14orf80, C5orf51, CDC37, CEP128, CUL2, DDIT3, ELAVL1, GSK3B, GSTP1, KIAA0101, MYH15, NCKAP5L, PABPC4L, PCNA, RBM27, RNF214, RPS6KA6, RSBN1, SCGB1D2, SLC38A10, SND1, TRIM28, UBC, VAV2, ZNF256, ZNF667, ZRANB3 | 27 | 16 | Cell Morphology, Cellular Assembly and Organization, Cellular Development |
| **4** | A2M, APOA1, APOB, APOC3, B3GNT9, chymotrypsin, Cytokeratin, elastase, FGA, FGB, FGG, Fibrin, Fibrinogen, GPIIB-IIIA, Growth hormone, HDL, HDL-cholesterol, HP, HPR, Kallikrein, KRT1, KRT9, KRT10, KRT6B, LDL-cholesterol, LRP, NFkB (complex), PCYOX1, PEBP1, Pro-inflammatory Cytokine, SAA, SFTPA1, SFTPD, Stat3-Stat3, VLDL-cholesterol | 27 | 18 | Developmental Disorder, Hematological Disease, Hereditary Disorder |
| **5** | ANXA3, APITD1, C1GALT1C1, C1orf86, C9orf72, CYC1, EIF2B2, EIF2B3, EMG1, FANCB, FANCE, FANCF, FANCM, GALNT2, GALNT5, GGA1, GGA3, KRT79, LANCL1, MON2, MYOZ1, NAPSA, NLE1, PNKP, RAB6B, RABGAP1, RAPGEF6, RBM34, RMI2, SH3BGRL, SLC25A24, STRA13, TOP3A, UBC, ZNF292 | 25 | 16 | Developmental Disorder, Hematological Disease, Hereditary Disorder |
| **6** | Alp, BMP2K, C16orf88, CBR1, CD3, CDC45, Cg, CHD3, CNGA2, CUL4B, DBI, ENO2, Focal adhesion kinase, Hdac, Histone h3, Histone h4, Hsp70, HSPA6, ICAM1, IDH1, IKK (complex), LDL, NADPH oxidase, P38 MAPK, Pdgf (complex), PI3K (complex), Pkc(s), PREX1, RNA polymerase II, Sod, SRC (family), TALDO1, Vegf, VNN1, WNT9B | 24 | 16 | Cancer, Gastrointestinal Disease, Cardiovascular Disease |
| **7** | AKAP6, BEND7, C11orf48, CCDC85A, CCNB1, CCND1, CDK5RAP3, CDKN1B, CMIP, DACH2, DDRGK1, DGCR14, DNAJC16, FOXO3, GSTM4, GSTM5, GSTO2, hemoglobin, LYAR, MYRIP, NANOG, PARPBP, PGAM4, PIK3R1, PRRC2C, RAB19, RSL24D1, SIX6, SLC34A2, STAT5A, TMEM55A, UBC, UBLCP1, UFC1, ZNF462 | 20 | 14 | Cardiovascular System Development and Function, Cell Cycle, Skeletal and Muscular System Development and Function |
| **8** | ADCYAP1, alcohol dehydrogenase, ALDH16A1, ALDOC, APP, ASXL2, C19orf40, CALML3, CASP6, CCL5, CRTAC1, CWF19L2, FBXO34, GSTM3, HSP90AB1, HSPA2, HSPB7, IRAK3, KIFC3, MDH1, NUCB1, PDE1A, PSMB4, PSMD1, PTMS, RAB10, RNASE1, RUSC1, SCAVENGER receptor CLASS A, SDCCAG8, SH3RF2, TAGLN2, TRAF6, USP1, ZBTB20 | 20 | 14 | Organismal Injury and Abnormalities, Cell Death and Survival, Nervous System Development and Function |
| **9** | B4GALT1, BCR (complex), Collagen(s), CTSZ, DPP4, ERK, Fc gamma receptor, GATA3, GOT1, HLA-C, HSP, Ifn, IFN Beta, Ifn gamma, IgG1, Igg3, IgG, IGHG2, Igm, Ikb, IKBKB, IL1, IL12 (complex), IL12 (family), Immunoglobulin, Interferon alpha, LGALS3BP, MHC Class I (complex), MHC CLASS I (family), MHC Class II (complex), NKX3-2, PPBP, PRSS8, Tgf beta, Tlr | 17 | 12 | Cellular Movement, Hematological System Development and Function, Immune Cell Trafficking |
| **10** | 26sProteasome, ADCY, ARHGAP24, Calmodulin, CD97, CELSR1, chemokine, Ck2, EMR2, endocannabinoid, FBP1, FSH, Gpcr, GPR4, GPR68, GPRC5A, GRM8, Insulin, MAP9, Mapk, MID2, NAPA, OCRL, Pka, PLC, Rac, RAPSN, Ras, Ras homolog, Sfk, Shc, SYTL4, Trk Receptor, UBE2N, Ubiquitin | 17 | 12 | Cellular Assembly and Organization, Cellular Function and Maintenance, Molecular Transport |
| **11** | ADH1B, AKAP9, ALDH3A1, Ap1, BLVRA, C2, C/ebp, calpain, caspase, Collagen type IV, Cyclin A, Cyclin E, DACH1, DLGAP4, estrogen receptor, FHL1, FN1, Hsp90, HTRA2, Integrin, Jnk, Lfa-1, MAP2K1/2, Mek, Metalloprotease, Mmp, MSLN, NFAT (complex), Nfat (family), NUMA1, p70 S6k, PDGF BB, PSMD14, TCR, trypsin | 16 | 13 | Cell Cycle, Visual System Development and Function, Hair and Skin Development and Function |
| **12** | ADRBK2, CACNA1B, CCM2, CHRM3, CNR1, COL11A2, COL2A1, CREB3L3, D-glucose, endocannabinoid, FCHSD2, GABBR1, GBP5, GNB4, GNG5, GNG7, GNGT1, GPM6A, GPR68, ITGB1BP1, JAKMIP1, KRIT1, N-type Calcium Channel, PLA2G6, PLCB3, RGS6, SEPT4, SEPT7, SEPT8, SH3BGR, TRHR, TRPV4, UNC13C, VCPIP1, ZNF219 | 13 | 10 | Connective Tissue Disorders, Developmental Disorder, Hereditary Disorder |
| **13** | ACP5, AKAP12, BCL3, CAMP, CPM, CSF1, CYP11A1, FANK1, FPR2, Hedgehog, HLX, HSD17B1, ITGB8, JUN, mannitol, MAP2K2, MAZ, MOGS, MTIF2, NOTCH4, PGC, PROM1, PTPRO, SERPINB2, SFTPB, SIRT6, SLC8A1, SMAD5, SOD2, STAB2, Stat3-Stat3, TEAD4, TMSB10/TMSB4X, USP36, VEGFA | 8 | 7 | Cardiovascular System Development and Function, Embryonic Development, Organismal Development |
| **14** | ANO8, COQ9 | 2 | 1 | Hereditary Disorder, Metabolic Disease, Cancer |
| **15** | Spag6, SPAG17 | 2 | 1 | Cellular Assembly and Organization, Cellular Compromise, Cellular Function and Maintenance |
| **16** | ADCY10, SLC9C1 | 2 | 1 | Cellular Movement, Reproductive System Development and Function, Reproductive System Disease |

- Red=upregulated proteins
- Green=downregulated protein

\# Networks = collections of interconnected molecules assembled by a network algorithm. Each connection represents known relationships between the molecules, found in the Ingenuity Knowledge Base.

* Score= The degree of relevance of Network Eligible molecules to the BALF dataset.  The score takes into account the number of Network Eligible molecules in the network and its size, as well as the total number of Network Eligible molecules analyzed and the total number of molecules in the Ingenuity Knowledge Base that could potentially be included in networks.  The network Score is based on the hypergeometric distribution and is calculated with the right-tailed Fisher's Exact Test. Score=-log(Fisher's Exact test result)

- ^ Focus Molecules= The number of proteins identified in the BALF proteome that is found in the network
