## Supplementary material for "Proteomic network analysis of bronchoalveolar lavage fluid in ex-smokers to discover implicated protein targets and novel drug treatments for chronic obstructive pulmonary disease": Table S1

- **Table S3: Unique proteins downregulated in BALF (n=139).** Differentially expressed proteins with at least 1.5x fold change decrease in the BALF proteome in COPD versus control cohort samples.

| - Uniprot ID | - Symbol | - Entrez Gene Name | - Location | - Type(s) | - Fold change | - Previously described association with COPD | - References |
| --- | --- | --- | --- | --- | --- | --- | --- |
| - Q9HCH0 | - NCKAP5L | - NCK-associated protein 5-like | - unknown | - other | - -8.55 | - NONE |  |
| - Q01995 | - TAGLN | - transgelin | - Cytoplasm | - other | - -8.11 | - NONE |  |
| - Q29865 | - HLA-C | - major histocompatibility complex, class I, C | - Plasma Membrane | - other | - -7.63 | - GWAS analysis in the ECLIPSE study noted a SNP in the HLA-C region | - ^53^ |
| - P14314 | - PRKCSH | - protein kinase C substrate 80K-H | - Cytoplasm | - enzyme | - -6.84 | - NONE |  |
| - B3KS81 | - SRRM5 | - serine/arginine repetitive matrix 5 | - unknown | - other | - -6.06 | - NONE |  |
| - P20142 | - PEPC | - Gastricsin | - Extracellular Space |  | - -5.25 | - NONE |  |
| - O95185 | - UNC5C | - unc-5 homolog C | - Plasma Membrane | - transmembrane receptor/ netrin | - -5.11 | - NONE |  |
| - Q9Y3P9 | - RABGAP1 | - RAB GTPase activating protein 1 | - Cytoplasm | - other | - -5.04 | - NONE |  |
| - Q7Z3U7 | - MON2 | - MON2 homolog (S. cerevisiae) | - Cytoplasm | - other | - -5.02 | - NONE |  |
| - Q9NWN3 | - FBXO34 | - F-box protein 34 | - unknown | - other | - -4.80 | - NONE |  |
| - O60885 | - BRD4 | - bromodomain containing 4 | - Nucleus | - kinase | - -4.33 | - NONE |  |
| - Q9UHX3 | - EMR2 | - egf-like module containing, Mucin-like, hormone receptor-like 2 | - Plasma Membrane | - G-protein coupled receptor | - -4.32 | - NONE |  |
| - Q9NSY1 | - BMP2K | - BMP2 inducible kinase | - Nucleus | - kinase | - -4.03 | - NONE |  |
| - Q9H0P7 | - CF059 | - Putative uncharacterized protein encoded by NCRNA00241 |  |  | - -3.96 | - NONE |  |
| - O75419 | - CDC45 | - cell division cycle 45 homolog | - Nucleus | - other | - -3.68 | - NONE |  |
| - P03950 | - ANG | - angiogenin, ribonuclease, RNase A family, 5 | - Extracellular Space | - enzyme | - -3.59 | - NONE |  |
| - Q8IWL2 | - SFTPA1 | - surfactant protein A1 | - Extracellular Space | - transporter | - -3.59 | - Imbalances of the surfactant proteins, major components of alveolar fluid have been implicated in COPD | - ^54-60^ |
| - Q1ED39 | - CP088 | - Protein C16orf88 |  |  | - -3.57 | - NONE |  |
| - P54750 | - PDE1A | - phosphodiesterase 1A, calmodulin-dependent | - Cytoplasm | - enzyme | - -3.48 | - NONE |  |
| - Q8ND24 | - RNF214 | - ring finger protein 214 | - unknown | - other | - -3.46 | - NONE |  |
| - Q96N16 | - JAKMIP1 | - janus kinase and microtubule interacting protein 1 | - Cytoplasm | - other | - -3.41 | - NONE |  |
| - Q14980 | - NUMA1 | - nuclear mitotic apparatus protein 1 | - Nucleus | - other | - -3.26 | - NONE |  |
| - Q9UI36 | - DACH1 | - dachshund homolog 1 | - Nucleus | - transcription regulator | - -3.03 | - NONE |  |
| - Q9UHG3 | - PCYOX1 | - prenylcysteine oxidase 1 | - Cytoplasm | - enzyme | - -3.03 | - NONE |  |
| - O14905 | - WNT9B | - wingless-type MMTV integration site family, member 9B | - Extracellular Space | - Signal transduction | - -3.03 | - NONE |  |
| - Q99996 | - AKAP9 | - A kinase (PRKA) anchor protein (yotiao) 9 | - Cytoplasm | - other | - -3.02 | - NONE |  |
| - Q9Y2P7 | - ZNF256 | - zinc finger protein 256 | - Nucleus | - transcription regulator | - -3.00 | - NONE |  |
| - P02747 | - C1QC | - complement component 1, q subcomponent, C chain | - Extracellular Space | - other | - -3.00 | - NONE |  |
| - O00750 | - PIK3C2B | - phosphoinositide-3-kinase, class 2, beta polypeptide | - Cytoplasm | - kinase | - -3.00 | - Associated with glucocorticoid sensitivity and inflammation in COPD | - ^61-63^ |
| - Q9P2N5 | - RBM27 | - RNA binding motif protein 27 | - Nucleus | - other | - -3.00 | - NONE |  |
| - Q9Y520 | - PRRC2C | - proline-rich coiled-coil 2C | - Cytoplasm | - other | - -2.92 | - NONE |  |
| - Q9P2Y4 | - ZNF219 | - zinc finger protein 219 | - Nucleus | - transcription regulator | - -2.92 | - NONE |  |
| - O43813 | - LANCL1 | - LanC lantibiotic synthetase component C-like 1 (bacterial) | - Plasma Membrane | - other | - -2.92 | - NONE |  |
| - P01714 | - LV301 | - Ig lambda chain V_III region SH | - Extracellular Space | - immunoglobin | - -2.92 | - NONE |  |
| - O75264 | - CS077 | - Transmembrane protein C19orf77 |  |  | - -2.92 | - NONE |  |
| - A6NMX2 | - EIF4E1B | - eukaryotic translation initiation factor 4E family member 1B | - unknown | - other | - -2.92 | - NONE |  |
| - Q8IYD8 | - FANCM | - Fanconi anemia, complementation group M | - Nucleus | - enzyme | - -2.92 | - NONE |  |
| - Q8TC84 | - FANK1 | - Fibronectin type III and ankyrin repeat domains 1 | - Nucleus | - transcription regulator | - -2.92 | - NONE |  |
| - Q96NX9 | - DACH2 | - dachshund homolog 2 | - Nucleus | - other | - -2.90 | - NONE |  |
| - Q9BVG8 | - KIFC3 | - kinesin family member C3 | - Cytoplasm | - enzyme | - -2.87 | - NONE |  |
| - O14920 | - IKBKB | - inhibitor of kappa light polypeptide gene enhancer in B-cells, kinase beta | - Cytoplasm | - kinase | - -2.75 | - Implicated in COPD inflammation | - ^64-66^ |
| - Q92738 | - USP6NL | - USP6 N-terminal like/ RAB5 effector RN-tre | - Plasma Membrane | - Cytoskeleton element involved in pinocytosis | - -2.75 | - NONE |  |
| - Q96JB5 | - CDK5RAP3 | - CDK5 regulatory subunit associated protein 3 | - Cytoplasm | - other | - -2.74 | - NONE |  |
| - Q9UEW3 | - MARCO | - macrophage receptor with collagenous structure | - Plasma Membrane | - transmembrane receptor | - -2.74 | - A macrophage scavenger receptor involved in bacterial phagocytosis in COPD | - ^67,68^ |
| - Q13724 | - MOGS | - mannosyl-oligosaccharide glucosidase | - Cytoplasm | - enzyme | - -2.68 | - NONE |  |
| - P51674 | - GPM6A | - glycoprotein M6A | - Plasma Membrane | - ion channel | - -2.68 | - NONE |  |
| - Q16651 | - PRSS8 | - protease, serine, 8 | - Extracellular Space | - peptidase | - -2.63 | - NONE |  |
| - O96009 | - NAPSA | - napsin A aspartic peptidase | - Extracellular Space | - peptidase | - -2.58 | - NONE |  |
| - Q9NVX2 | - NLE1 | - notchless homolog 1 (Drosophila) | - Nucleus | - enzyme | - -2.54 | - NONE |  |
| - P02751 | - FN1 | - fibronectin 1 | - Extracellular Space | - enzyme | - -2.53 | - Matrix protein involved in fibroblast proliferation implicated in COPD pathogenesis | - ^38,69-73^ |
| - Q13023 | - AKAP6 | - A kinase (PRKA) anchor protein 6 | - Nucleus | - other | - -2.53 | - NONE |  |
| - Q5VWQ0 | - RSBN1 | - round spermatid basic protein 1 | - Nucleus | - other | - -2.53 | - NONE |  |
| - Q9UGM5 | - FETUB | - Fetuin B | - Extracellular Space | - other | - -2.49 | - NONE |  |
| - Q9Y2G8 | - DNAJC16 | - DnaJ (Hsp40) homolog, subfamily C, member 16 | - unknown | - other | - -2.45 | - NONE |  |
| - P35247 | - SFTPD | - surfactant protein D | - Extracellular Space | - other | - -2.44 | - Imbalances of the surfactant proteins, major components of alveolar fluid have been implicated in COPD | - ^54,56,74-79^ |
| - Q2TBE0 | - CWF19L2 | - CWF19-like 2, cell cycle control | - unknown | - other | - -2.42 | - NONE |  |
| - Q9BYF1 | - ACE2 | - angiotensin I converting enzyme (peptidyl-dipeptidase A) 2 | - Plasma Membrane | - peptidase | - -2.41 |  | - ^80^ |
| - O95969 | - SCGB1D2 | - secretoglobin, family 1D, member 2 | - Extracellular Space | - other | - -2.40 | - NONE |  |
| - P78367 | - NKX32 | - Homeobox protein Nkx_3.2 |  |  | - -2.40 | - NONE |  |
| - Q9P275 | - USP36 | - ubiquitin specific peptidase 36 | - Nucleus | - peptidase | - -2.40 | - NONE |  |
| - O43464 | - HTRA2 | - HtrA serine peptidase 2 | - Cytoplasm | - peptidase | - -2.40 | - NONE |  |
| - O60281 | - ZNF292 | - zinc finger protein 292 | - Nucleus | - transcription regulator | - -2.38 | - NONE |  |
| - Q96JM2 | - ZNF462 | - zinc finger protein 462 | - Nucleus | - other | - -2.37 | - NONE |  |
| - P27487 | - DPP4 | - dipeptidyl-peptidase 4 | - Plasma Membrane | - peptidase | - -2.33 | - Putative serum COPD biomarker | - ^81^ |
| - Q86SX3 | - CN080 | - Uncharacterized protein C14orf80 |  |  | - -2.28 | - NONE |  |
| - Q9UJV3 | - MID2 | - midline 2 | - Cytoplasm | - other | - -2.13 | - NONE |  |
| - Q6ZU80 | - CEP128 | - centrosomal protein 128kDa | - unknown | - other | - -2.12 | - NONE |  |
| - Q01968 | - OCRL | - Inositol polyphosphate 5_phosphatase | - Cytoplasm | - phosphatase | - -2.11 | - NONE |  |
| - P42696 | - RBM34 | - RNA binding motif protein 34 | - Nucleus | - other | - -2.11 | - NONE |  |
| - P0CB38 | - PABPC4L | - poly(A) binding protein, cytoplasmic 4-like | - unknown | - other | - -2.09 | - NONE |  |
| - P49770 | - EIF2B2 | - eukaryotic translation initiation factor 2B, subunit 2 beta, 39kDa | - Cytoplasm | - translation regulator | - -2.09 | - NONE |  |
| - P11678 | - EPX | - eosinophil peroxidase | - Cytoplasm | - enzyme | - -2.06 | - NONE, Although one reference examined EPX in subjects, it is was NOT differentially expressed in COPD | - ^82^ |
| - B1AJZ9 | - FHAD1 | - forkhead-associated (FHA) phosphopeptide binding domain 1 | - unknown | - other | - -2.04 | - NONE |  |
| - P54920 | - NAPA | - N-ethylmaleimide-sensitive factor attachment protein, alpha | - Cytoplasm | - other | - -2.03 | - NONE |  |
| - P01771 | - HV310 | - Ig heavy chain V_III region HIL | - Extracellular Space | - immunoglobin | - -2.03 | - NONE |  |
| - Q9Y2H0 | - DLGAP4 | - discs, large homolog-associated protein 4 | - Plasma Membrane | - other | - -2.01 | - NONE |  |
| - Q07954 | - LRP1 (includes EG:16971) | - low density lipoprotein receptor-related protein 1 | - Plasma Membrane | - transmembrane receptor | - -2.00 | - NONE |  |
| - Q13702 | - RAPSN | - receptor-associated protein of the synapse | - Plasma Membrane | - other | - -2.00 | - NONE |  |
| - Q13620 | - CUL4B | - cullin 4B | - Nucleus | - other | - -2.00 | - NONE |  |
| - P15291 | - B4GALT1 | - UDP-Gal:betaGlcNAc beta 1,4- galactosyltransferase, polypeptide 1 | - Cytoplasm | - enzyme | - -1.99 | - NONE |  |
| - P04259 | - K2C6B | - Keratin type II cytoskeletal 6B |  |  | - -1.99 | - NONE |  |
| - P14384 | - CPM | - carboxypeptidase M | - Plasma Membrane | - peptidase | - -1.98 | - NONE |  |
| - Q5HYK9 | - ZNF667 | - zinc finger protein 667 | - Nucleus | - other | - -1.97 | - NONE |  |
| - Q5VVM6 | - CCDC30 | - coiled-coil domain containing 30 | - unknown | - other | - -1.97 | - NONE |  |
| - Q16181 | - SEP7 | - septin 7 | - Cytoplasm | - other | - -1.97 | - NONE |  |
| - P01763 | - HV302 | - Ig heavy chain V_III region WEA | - Extracellular Space | - immunoglobin | - -1.96 | - NONE |  |
| - P07225 | - PROS1 | - protein S (alpha) | - Extracellular Space | - other | - -1.96 | - NONE |  |
| - P01780 | - HV319 | - Ig heavy chain V_III region JON | - Extracellular Space | - immunoglobin | - -1.95 | - NONE |  |
| - P01605 | - KV113 | - Ig kappa chain V_I region Lay | - Extracellular Space | - immunoglobin | - -1.95 | - NONE |  |
| - P01611 | - KV119 | - Ig kappa chain V_I region Wes | - Extracellular Space | - immunoglobin | - -1.93 | - NONE |  |
| - P01612 | - KV120 | - Ig kappa chain V_I region Mev | - Extracellular Space | - immunoglobin | - -1.93 | - NONE |  |
| - Q8NB66 | - UNC13C | - unc-13 homolog C | - Cytoplasm | - other | - -1.90 | - NONE |  |
| - A4D1S5 | - RAB19 | - RAB19, member RAS oncogene family | - Cytoplasm | - enzyme | - -1.90 | - NONE |  |
| - Q9Y613 | - FHOD1 | - formin homology 2 domain containing 1 | - Nucleus | - other | - -1.89 | - NONE |  |
| - Q86Y33 | - CDC20B | - cell division cycle 20 homolog B | - unknown | - other | - -1.87 | - NONE |  |
| - P06317 | - LV603 | - Ig lambda chain V_VI region SUT | - Extracellular Space | - immunoglobin | - -1.86 | - NONE |  |
| - Q5FWF4 | - ZRANB3 | - zinc finger, RAN-binding domain containing 3 | - unknown | - enzyme | - -1.83 | - NONE |  |
| - Q0VAM2 | - RASGEF1B | - RasGEF domain family, member 1B | - unknown | - other | - -1.82 | - NONE |  |
| - Q9HAV0 | - GNB4 | - guanine nucleotide binding protein (G protein), beta polypeptide 4 | - Plasma Membrane | - enzyme | - -1.79 | - NONE |  |
| - P16070 | - CD44 | - CD44 molecule (Indian blood group) | - Plasma Membrane | - enzyme | - -1.79 | - CD44 HA receptor is implicated in macrophage phagocytic ability and appears to be decreased in COPD | - ^9,83-86^ |
| - P16144 | - ITGB4 | - integrin, beta 4 | - Plasma Membrane | - transmembrane receptor | - -1.77 | - NONE |  |
| - Q12873 | - CHD3 | - chromodomain helicase DNA binding protein 3 | - Nucleus | - enzyme | - -1.75 | - NONE |  |
| - P15151 | - PVR | - poliovirus receptor | - Plasma Membrane | - other | - -1.75 | - NONE |  |
| - Q08380 | - LGALS3BP | - lectin, galactoside-binding, soluble, 3 binding protein | - Plasma Membrane | - transmembrane receptor | - -1.75 | - NONE for the specific protein, but as a binding protein for a lectin, its associates with galectin 3 which seems to regulate macrophage efferocytosis in COPD, , | - ^26,27^ |
| - Q9HBR0 | - SLC38A10 | - solute carrier family 38, member 10 | - unknown | - other | - -1.74 | - NONE |  |
| - Q6Q759 | - SPAG17 | - sperm associated antigen 17 | - unknown | - other | - -1.73 | - NONE |  |
| - Q15582 | - TGFBI | - transforming growth factor, beta-induced, 68kDa | - Extracellular Space | - other | - -1.73 | - NONE |  |
| - A6NCL7 | - ANKRD33B | - ankyrin repeat domain 33B | - unknown | - other | - -1.72 | - NONE |  |
| - Q8TEU7 | - RAPGEF6 | - Rap guanine nucleotide exchange factor (GEF) 6 | - Plasma Membrane | - other | - -1.72 | - NONE |  |
| - P01619 | - KV301 | - Ig kappa chain V_III region B6 | - Extracellular Space | - immunoglobin | - -1.72 | - NONE |  |
| - Q8TCU6 | - PREX1 | - phosphatidylinositol-3,4,5-trisphosphate-dependent Rac exchange factor 1 | - Cytoplasm | - other | - -1.70 | - NONE |  |
| - P04114 | - APOB | - apolipoprotein B (including Ag(x) antigen) | - Extracellular Space | - Lipid metabolism | - -1.68 | - NONE |  |
| - P06314 | - KV40 | - Ig kappa chain V_IV region B17 | - Extracellular Space | - immunoglobin | - -1.67 | - NONE |  |
| - P01859 | - IGHG2 | - immunoglobulin heavy constant gamma 2 (G2m marker) | - Plasma Membrane | - immunoglobin | - -1.67 | - NONE |  |
| - P17174 | - GOT1 | - glutamic-oxaloacetic transaminase 1, soluble (aspartate aminotransferase 1) | - Cytoplasm | - enzyme | - -1.67 | - NONE |  |
| - Q02818 | - NUCB1 | - nucleobindin 1 | - Cytoplasm | - other | - -1.67 | - NONE |  |
| - P05362 | - ICAM1 | - intercellular adhesion molecule 1 | - Plasma Membrane | - transmembrane receptor | - -1.66 | - Discordant of ICAM in COPD in literature compared to our findings | - ^87-96^ |
| - Q6UX72 | - B3GNT9 | - UDP-GlcNAc:betaGal beta-1,3-N-acetylglucosaminyltransferase 9 | - unknown | - enzyme | - -1.65 | - NONE |  |
| - P52823 | - STC1 | - stanniocalcin 1 | - Extracellular Space | - kinase | - -1.65 | - NONE |  |
| - P36222 | - CHI3L1 | - chitinase 3-like 1 | - Extracellular Space | - Tissue remodeling | - -1.65 | - Increased in the serum and BAL of smokers with COPD compared to never smokers or smokers without COPD | - ^97-99^ |
| - Q96JH7 | - VCPIP1 | - valosin containing protein (p97)/p47 complex interacting protein 1 | - Cytoplasm | - peptidase | - -1.64 | - NONE |  |
| - O95436 | - SLC34A2 | - solute carrier family 34 (sodium phosphate), member 2 | - Plasma Membrane | - transporter | - -1.64 | - NONE |  |
| - P46199 | - MTIF2 | - mitochondrial translational initiation factor 2 | - Cytoplasm | - translation regulator | - -1.64 | - NONE |  |
| - Q86SQ7 | - SDCCAG8 | - serologically defined colon cancer antigen 8 | - Cytoplasm | - other | - -1.64 | - NONE |  |
| - Q8N7W2 | - BEND7 | - BEN domain containing 7 | - unknown | - other | - -1.61 | - NONE |  |
| - P01024 | - C3 | - complement component 3 | - Extracellular Space | - peptidase | - -1.58 | - Serum levels of C3 was decreased in serum and sputum of subjects with COPD | - ^100-102^ |
| - P07998 | - RNASE1 | - ribonuclease, RNase A family, 1 (pancreatic) | - Extracellular Space | - enzyme | - -1.58 | - NONE |  |
| - P00739 | - HPR | - haptoglobin-related protein | - Extracellular Space | - peptidase | - -1.58 | - NONE |  |
| - Q76L83 | - ASXL2 | - additional sex combs like 2 | - Extracellular Space | - other | - -1.57 | - NONE |  |
| - Q8NFJ5 | - GPRC5A | - G protein-coupled receptor, family C, group 5, member A | - Plasma Membrane | - G-protein coupled receptor | - -1.57 | - Decreased in lung epithelia associated with lung adenocarcinoma compared to epithelia from subjects with COPD or never smokers | - ^103,104^ |
| - Q7Z7M9 | - GALNT5 | - UDP-N-acetyl-alpha-D-galactosamine:polypeptide N-acetylgalactosaminyltransferase 5 (GalNAc-T5) | - Cytoplasm | - enzyme | - -1.57 | - NONE |  |
| - P06331 | - HV209 | - Ig heavy chain V_II region ARH_77 | - Extracellular Space | - immunoglobin | - -1.57 | - NONE |  |
| - P15941 | - MUC1 | - Mucin 1, cell surface associated | - Plasma Membrane | - transcription regulator | - -1.56 | - Levels are affected by age and smoking in lung tissue, sputum and plasma | - ^56,105,106^ |
| - P48960 | - CD97 | - CD97 molecule | - Plasma Membrane | - G-protein coupled receptor | - -1.55 | - NONE |  |
| - P06681 | - C2 | - complement component 2 | - Extracellular Space | - peptidase | - -1.54 | - NONE |  |
| - Q9NYQ6 | - CELSR1 | - cadherin, EGF LAG seven-pass G-type receptor 1 | - Plasma Membrane | - G-protein coupled receptor | - -1.53 | - NONE |  |
| - P0CG06 | - IGLC3 | - immunoglobulin lambda constant 3 (Kern-Oz+ marker) | - Extracellular Space | - other | - -1.53 | - NONE |  |
| - P06126 | - CD1A | - CD1a molecule | - Plasma Membrane | - other | - -1.52 |  | - ^107-110^ |
| - O00560 | - SDCBP | - syndecan binding protein (syntenin) | - Plasma Membrane | - enzyme | - -1.51 | - NONE |  |
