## Supplementary material for "Proteomic network analysis of bronchoalveolar lavage fluid in ex-smokers to discover implicated protein targets and novel drug treatments for chronic obstructive pulmonary disease": Table S2

- **Table S2:**  **Unique proteins upregulated in BALF (n=95),** Differentially expressed proteins with at least 1.5x fold change increase in the BALF proteome in COPD versus control cohort samples.

| Uniprot ID | Symbol | Entrez Gene Name | Location | Type(s) | Fold change | Previously described association with COPD | References |
| --- | --- | --- | --- | --- | --- | --- | --- |
| P00325 | ADH1B | alcohol dehydrogenase 1B (class I), beta polypeptide | Cytoplasm | Lipid metabolism | 12.86 | NONE |  |
| P46439 | GSTM5 | glutathione S-transferase mu 5 | Cytoplasm | enzyme | 11.26 | Various GST polymorphisms implicated in lung protection | ^3-5^ |
| O00487 | PSMD14 | proteasome (prosome, macropain) 26S subunit, non-ATPase, 14 | Cytoplasm | peptidase | 9.53 | NONE |  |
| Q86TZ1 | TTC6 | tetratricopeptide repeat domain 6 | unknown | other | 8.68 | NONE |  |
| P02743 | APCS | amyloid P component, serum | Extracellular Space | pentraxin | 8.23 | Not APCS but Pentraxin 3 in COPD | ^6-8^ |
| Q9UBR2 | CTSZ | cathepsin Z | Cytoplasm | peptidase | 7.92 | NONE |  |
| Q9NP98 | MYOZ1 | myozenin 1 | Cytoplasm | other | 7.08 | NONE |  |
| Q13402 | MYO7A | myosin VIIA | Cytoplasm | enzyme | 6.84 | NONE |  |
| Q49MG5 | MAP9 | microtubule-associated protein 9 | unknown | other | 6.31 | NONE |  |
| Q13642 | FHL1 | four and a half LIM domains 1 | Cytoplasm | other | 5.53 | NONE |  |
| P02775 | PPBP | pro-platelet basic protein (chemokine (C-X-C motif) ligand 7) | Extracellular Space | cytokine | 5.47 | Neutrophil marker increased in severe stable COPD | ^9^ |
| P21333 | FLNA | filamin A, alpha | Cytoplasm | other | 5.14 | NONE |  |
| Q8NFC6 | BOD1L1 | biorientation of chromosomes in cell division 1-like 1 | Extracellular Space | other | 4.43 | NONE |  |
| P04220 | MUCB | Ig mu heavy chain disease protein | Plasma Membrane |  | 4.36 | NONE |  |
| P02656 | APOC3 | apolipoprotein C-III | Extracellular Space | Lipid metabolism | 4.24 | NONE |  |
| P42330 | AKR1C3 | aldo-keto reductase family 1, member C3 (3-alpha hydroxysteroid dehydrogenase, type II) | Cytoplasm | enzyme | 4.11 | NONE |  |
| P35749 | MYH11 | myosin, heavy chain 11, smooth muscle | Cytoplasm | other | 4.01 | Myosin heavy chain variation was noted in COPD but not specific proteins | ^10-12^ |
| O00522 | KRIT1 | KRIT1, ankyrin repeat containing | Plasma Membrane | other | 3.89 | NONE |  |
| A6NDU8 | CE051 | UPF0600 protein C5orf51 | unknown | other | 3.56 | NONE |  |
| P04114 | APOB | apolipoprotein B (including Ag(x) antigen) | Extracellular Space | transporter | 3.556 | NONE |  |
| Q5XKE5 | KRT79 | keratin 79 | Extracellular Space | other | 3.43 | NONE |  |
| Q9NWS1 | PARPBP | PARP1 binding protein | Nucleus | other | 3.42 | NONE |  |
| P12429 | ANXA3 | Annexin A3 |  |  | 3.34 | NONE |  |
| P30838 | ALDH3A1 | aldehyde dehydrogenase 3 family, member A1 | Cytoplasm | Lipid metabolism | 3.33 | None, but is implicated in cell proliferation |  |
| Q6NUK1 | SLC25A24 | solute carrier family 25 (mitochondrial carrier; phosphate carrier), member 24 | Cytoplasm | other | 3.33 | NONE |  |
| P00738 | HP | haptoglobin | Extracellular Space | peptidase | 3.22 | Acute phase reactant associated with COPD | ^13,14^ |
| Q16787 | LAMA3 | laminin, alpha 3 | Extracellular Space | other | 3.2 | Haemophilus and Moraxella binds to laminin |  |
| P02675 | FGB | fibrinogen beta chain | Extracellular Space | other | 3.17 | Serum fibrinogen levels in COPD associated with exacerbations | ^14-20^ |
| P08670 | VIM | vimentin | Cytoplasm | cytoskeleton component | 3.12 | Epithelial to mesenchymal transition | ^21-25^ |
| Q7KZI7 | MARK2 | MAP/microtubule affinity-regulating kinase 2 | Cytoplasm | kinase | 3.12 | NONE |  |
| Q15847 | APM2 | Adipose most abundant gene transcript 2 protein |  |  | 3.01 | NONE |  |
| Q16280 | CNGA2 | cyclic nucleotide gated channel alpha 2 | Plasma Membrane | ion channel | 3.01 | NONE |  |
| O94782 | USP1 | ubiquitin specific peptidase 1 | Cytoplasm | peptidase | 2.9 | NONE |  |
| Q96C24 | SYTL4 | synaptotagmin-like 4 | Cytoplasm | transporter | 2.82 | NONE |  |
| P02671 | FIBA | Fibrinogen alpha chain |  |  | 2.78 | Serum fibrinogen levels in COPD associated with exacerbations | ^14-20^ |
| P02679 | FGG | fibrinogen gamma chain | Extracellular Space | other | 2.75 | Serum fibrinogen levels in COPD associated with exacerbations | ^14-20^ |
| P53004 | BLVRA | biliverdin reductase A | Cytoplasm | enzyme | 2.66 | NONE |  |
| Q2TVT3 | KGFLP2 | keratinocyte growth factor-like protein 2 | unknown | other | 2.62 | NONE |  |
| P23284 | PPIB | peptidylprolyl isomerase B (cyclophilin B) | Cytoplasm | enzyme | 2.55 | NONE |  |
| P17931 | LGALS3 | lectin, galactoside-binding, soluble, 3 | Extracellular Space | other | 2.52 | Increased Gal-3 in small airways | ^26,27^ |
| P13645 | K1C10 | Keratin type I cytoskeletal 10 |  |  | 2.47 | NONE |  |
| Q9NX58 | LYAR | Ly1 antibody reactive homolog (mouse) | Plasma Membrane | other | 2.37 | NONE |  |
| O95497 | VNN1 | vanin 1 | Plasma Membrane | Enzyme/lipid metabolism | 2.34 | NONE |  |
| P16152 | CBR1 | carbonyl reductase 1 | Cytoplasm | enzyme | 2.34 | NONE |  |
| P23771 | GATA3 | GATA binding protein 3 | Nucleus | transcription regulator | 2.33 | NONE |  |
| P01023 | A2M | alpha-2-macroglobulin | Extracellular Space | transporter | 2.31 | A protease inhibitor that has increased serum levels found in patients with alpha-1 antitrypsin deficiency | ^13,28-31^ |
| Q15149 | PLEC | plectin | Cytoplasm | other | 2.31 | NONE |  |
| P62328 | TYB6 | Thymosin beta_4 |  |  | 2.28 | NONE |  |
| Q6UXR4 | SERPINA13 | serpin peptidase inhibitor, clade A (alpha-1 antiproteinase, antitrypsin), member 13 (pseudogene) | Extracellular Space | other | 2.26 | A serpin peptidase inhibitor that is in the same family of peptidase inhibitor as alpha-1 antitrypsin (a serpin peptidase inhibitor, clade A, member 1) implicated in protease-antiprotease homeostasis | ^32,33^ |
| Q14019 | COTL1 | coactosin-like 1 | Cytoplasm | other | 2.25 | NONE |  |
| P03950 | ANGI | Angiogenin |  |  | 2.24 | Increased in induced sputum from stable COPD individuals compared to healthy smokers | ^34^ |
| Q9UK76 | HN1 | hematological and neurological expressed 1 | Nucleus | other | 2.21 | NONE |  |
| P02647 | APOA1 | apolipoprotein A-I | Extracellular Space | Lipid metabolism | 2.13 | COPD biomarker | ^35^ |
| P07108 | DBI | diazepam binding inhibitor (GABA receptor modulator, acyl-CoA binding protein) | Cytoplasm | other | 2.13 | NONE |  |
| P55822 | SH3BGR | SH3 domain binding glutamic acid-rich protein | Cytoplasm | other | 2.13 | NONE |  |
| P08758 | ANXA5 | annexin A5 | Plasma Membrane | Apoptosis pathway | 2.1 | Decreases macrophage efferocytosis and elastase-induced pulmonary emphysema in mice | ^36^ |
| P37837 | TALDO1 | transaldolase 1 | Cytoplasm | enzyme | 2.09 | NONE |  |
| P04259 | KRT6B | keratin 6B | Cytoplasm | other | 2.049 | NONE |  |
| P41222 | PTGDS | prostaglandin D2 synthase 21kDa (brain) | Cytoplasm | enzyme | 2.03 | Increased RNA expression in the human lung tissue of subjects with moderate versus mild COPD | ^37^ |
| Q9BWM5 | ZNF416 | zinc finger protein 416 | Nucleus | other | 1.98 | NONE |  |
| Q9HCE9 | ANO8 | anoctamin 8 | Extracellular Space | other | 1.98 | NONE |  |
| Q96PP8 | GBP5 | guanylate binding protein 5 | Plasma Membrane | enzyme | 1.95 | NONE |  |
| Q92888 | ARHGEF1 | Rho guanine nucleotide exchange factor (GEF) 1 | Cytoplasm | other | 1.94 | NONE |  |
| P51884 | LUM | lumican | Extracellular Space | other | 1.93 | Extracellular matrix component | ^38^ |
| P62937 | PPIA | Peptidyl_prolyl cis_trans isomerase A |  |  | 1.92 | Increased in lung tissue from smokers with COPD versus never-smokers, and non-COPD smokers | ^39^ |
| P09972 | ALDOC | aldolase C, fructose-bisphosphate | Cytoplasm | Metabolic enzyme | 1.91 | NONE |  |
| Q5JYT7 | KIAA1755 | KIAA1755 | unknown | other | 1.91 | NONE |  |
| P30086 | PEBP1 | phosphatidylethanolamine binding protein 1 | Cytoplasm | other | 1.9 | NONE |  |
| O75368 | SH3BGRL | SH3 domain binding glutamic acid-rich protein like | Cytoplasm | other | 1.89 | NONE |  |
| Q4G0N8 | SLC9C1 | solute carrier family 9, subfamily C (Na+-transporting carboxylic acid decarboxylase), member 1 | unknown | other | 1.89 | NONE |  |
| O75874 | IDH1 | isocitrate dehydrogenase 1 (NADP+), soluble | Cytoplasm | enzyme | 1.88 | NONE |  |
| Q13421 | MSLN | mesothelin | Extracellular Space | other | 1.88 | NONE |  |
| Q9Y6W5 | WASF2 | WAS protein family, member 2 | Cytoplasm | cytoskeleton | 1.87 | NONE |  |
| P50224 | ST1A3 | Sulfotransferase 1A3/1A4 |  |  | 1.86 | NONE |  |
| Q9Y2K3 | MYH15 | myosin, heavy chain 15 | Extracellular Space | other | 1.86 | Muscle dysfunction and aberrations of myosin composition within muscle has been associated with COPD | ^10,11,40-45^ |
| Q16881 | TXNRD1 | thioredoxin reductase 1 | Cytoplasm | enzyme | 1.82 | NONE |  |
| P37802 | TAGLN2 | transgelin 2 | Cytoplasm | other | 1.73 | NONE |  |
| P35527 | KRT9 | keratin 9 | Cytoplasm | other | 1.71 | NONE |  |
| P09104 | ENO2 | enolase 2 (gamma, neuronal) | Cytoplasm | enzyme | 1.7 | NONE |  |
| P40925 | MDH1 | malate dehydrogenase 1, NAD (soluble) | Cytoplasm | enzyme | 1.68 | NONE |  |
| P30041 | PRDX6 | peroxiredoxin 6 | Cytoplasm | enzyme | 1.66 | NONE |  |
| P04264 | K2C1 | Keratin type II cytoskeletal 1 |  |  | 1.65 | NONE |  |
| P61088 | UBE2N | ubiquitin-conjugating enzyme E2N | Cytoplasm | enzyme | 1.65 | CS induces UBE2N | ^46^ |
| P06319 | LV605 | Ig lambda chain V_VI region EB4 | Extracellular Space | immunoglobin | 1.64 | NONE |  |
| P20962 | PTMS | parathymosin | Nucleus | other | 1.63 | NONE |  |
| Q8N0Y7 | PGAM4 | phosphoglycerate mutase family member 4 | unknown | phosphatase | 1.63 | NONE |  |
| P06733 | ENO1 | enolase 1, (alpha) | Cytoplasm | transcription regulator | 1.61 | NONE |  |
| P09467 | FBP1 | fructose-1,6-bisphosphatase 1 | Cytoplasm | phosphatase | 1.6 | NONE |  |
| P17066 | HSPA6 | heat shock 70kDa protein 6 (HSP70B') | unknown | other | 1.59 | Increased proteins levels in patients with COPD treated with Inhaled Corticosteroids | ^47-49^ |
| Q96PX6 | CCDC85A | coiled-coil domain containing 85A | unknown | other | 1.57 | NONE |  |
| P23528 | CFL1 | cofilin 1 (non-muscle) | Nucleus | cytoskeleton | 1.56 | NONE |  |
| P63261 | ACTG | Actin_ cytoplasmic 2 | Cytoplasm |  | 1.56 | NONE |  |
| P06703 | S100A6 | S100 calcium binding protein A6 | Cytoplasm | transporter | 1.55 | Calcium binding protein involved in neutrophil activation and protein levels elevated in sputum from COPD versus control subjects | ^50-52^ |
