## Supplementary material for "Proteomic network analysis of bronchoalveolar lavage fluid in ex-smokers to discover implicated protein targets and novel drug treatments for chronic obstructive pulmonary disease": Table S1

**Table S1: Drugs commonly prescribed for stable COPD.** Curated drugs currently indicated for COPD ("MESH:D029424") from the Comparative Toxicogenomics Database^2^ and manually curated to include drugs used to treat stable COPD in the United States.

beclomethasone dipropionate

budesonide

arformoterol

roflumilast

aclidinium

fluticasone propionate

dexamethasone acetate

prednisolone tebutate

prednisone acetate

prednisolone acetate

mometasone

fluticasone

methyl prednisone

pirbuterol

ciclesonide

erdosteine

terbutaline

fluticasone furoate

glycopyrronium

indacaterol

ipratropium

levosalbutamol

clarithromycin

olodaterol

salbutamol

salmeterol

revefenacin

theophylline

salbutamol

aminophylline

umeclidinium

vilanterol

prednisone

tiotropium
