## Supplementary methods for "Proteomic network analysis of bronchoalveolar lavage fluid in ex-smokers to discover implicated protein targets and novel drug treatments for chronic obstructive pulmonary disease"

- **Supplementary methods**
- **Inclusion/exclusion criteria^1^**

For the COPD group, exclusion criteria included a forced expiratory volume at 1 s (FEV_1_) of <35%, hypercapnia, and comorbid diseases that would render bronchoscopy unsafe. Inclusion criteria were (1) chronic bronchitis as determined by history and/or emphysema as determined by chest x-ray or computed tomography; (2)  20 pack-years of cigarette smoking but smoking cessation for at least 1 year preceding enrollment; (3) the absence of other lung disease, including asthma and bronchiectasis; (4) chest x-ray findings that were normal or compatible with COPD but no other disease detected; (5) FEV_1:_forced vital capacity ratio and FEV_1_ both below the lower 95% confidence limit of normal on spirometry; (6) no atopy in the medical history; (7) <15% bronchodilator response with inhaled albuterol on spirometry; and (8) no antibiotic or steroid use for 4 weeks preceding enrollment. Inclusion criteria for the no-COPD group were the same as those for the COPD group, except for the absence of lung disease by clinical evaluation, normal chest x-ray, and normal spirometric results. Healthy nonsmokers met all inclusion criteria for the no-COPD group, except that all had <5 cumulative pack-years of smoking.
